# Self-immunity towards a novel competence-induced streptococcal murein hydrolase is mediated by a Fem-transferase-like protein

**DOI:** 10.1101/2024.09.27.615381

**Authors:** Marita Torrissen Mårli, Magnus Øverlie Arntzen, Jennie Ann Allred, Anna Teigen Schultheiss, Oddvar Oppegaard, Morten Kjos, Daniel Straume

**Author notes:** These authors contributed equally (Morten Kjos, Daniel Straume). Correspondence to: Morten Kjos or Daniel Straume.

## Abstract

Murein hydrolases (or peptidoglycan hydrolases) play diverse roles in bacteria, from controlled remodeling of the bacterial cell wall to lytic agents. In streptococci, a subset of these hydrolases is associated with competence-induced fratricide, a process where bacteria kill closely related cells to release DNA that can be taken up during natural transformation. Here, we characterize ScrM, a competence-induced murein hydrolase from *Streptococcus dysgalactiae* comprising a CHAP domain, an SH3b domain and an uncharacterized C-terminal domain (CCD). ScrM displayed lytic activity against pyogenic and salivarius group streptococci. Microscopy analysis of fluorescent fusions revealed that ScrM specifically localizes to the division zone of sensitive cells, with binding and localization mediated primarily by CCD. Upon competence induction, cells became immune to ScrM due to expression of ScrI, a Fem-transferase-like protein. We show by LC-MS/MS that ScrI incorporates Thr in place of Ala into the interpeptide bridges of peptidoglycan, which in turn prevents ScrM binding to the division zone, thereby protecting the cells from self-lysis during competence. ScrM and ScrI are conserved among pyogenic streptococcal pathogens and represent new players in the cell wall biogenesis of these bacteria that may form a platform for development of novel antimicrobial strategies.

## Introduction

The primary component of the bacterial cell wall, peptidoglycan (PG), is a polymeric mesh-like structure consisting of peptide-linked glycan strands which is required for bacteria to maintain their structure and morphology. A large diversity of enzymes targeting PG, collectively known as murein hydrolases (or PG hydrolases), are found across various biological systems in nature. Murein hydrolases include enzymes that are involved in the tightly regulated expansion and remodeling of PG during cell division, as well as in the separation of daughter cells (1). Additionally, some murein hydrolases degrade PG in an uncontrolled manner, leading to cell lysis, such as those produced by lytic phages, by bacteria undergoing programmed cell death, by eukaryotic innate immune systems, or by bacteria that uses murein hydrolases to lyse closely related competitors (1–4). When bacteria use murein hydrolases for competition, the producer cells need to protect themselves from self-killing through dedicated immunity mechanisms. However, the exact mechanisms underlying such self-immunity and the interplay between murein hydrolases and their corresponding immunity systems often remain obscure.

PG is composed of glycan strands made of alternating β-1,4-linked N-acetylglucosamine (GlcNAc) and N-acetylmuramic acid (MurNAc), which are crosslinked via pentapeptides bound to MurNAc. In most Gram-positive bacteria, including streptococci, the pentapeptide typically consists of L-Ala, D-Glu or D-iGln, L-Lys, and two D-Ala residues (5). Crosslinking occurs between the ε amino group of L-Lys of one acceptor pentapeptide and the carboxyl group of D-Ala in fourth position of a donor pentapeptide, a reaction powered by the release of D-Ala from the fifth position of the donor peptide (5–7). In many instances, additional amino acids are coupled to the ε amino group of L-lysine, resulting in a branched pentapeptide prior to interpeptide crosslinking (5, 7). Addition of such amino acids is catalyzed by Fem-transferases, non-ribosomal peptidyl transferases that sequentially add one or more amino acids to L-lysine of the lipid II peptidoglycan precursor (8). The resulting extra spacer in the peptide crosslink, commonly referred to as interpeptide bridges, varies in composition and length across different bacterial species and has been fully characterized for only a subset of them. In the human pathogen *Streptococcus pneumoniae,* the interpeptide bridge between the L-Lys and D-Ala of two crosslinked stem peptides consists of either Ser-Ala or Ala-Ala. The first L-Ala or L-Ser is added by the enzyme MurM and the second L-Ala by MurN (5, 9). For other streptococci, the interpeptide bridges typically include variations of this, such as Ala-Ala-(Ala) for the pathogenic *Streptococcus pyogenes* and *Streptococcus dysgalactiae* and Thr-Ala or Thr-Ser in *Streptococcus mutans* (7) (Table S1).

Murein hydrolases target different bonds within the PG structure, and are broadly classified into three groups based on the specific bonds they cleave: (i) glycosidases cleave glycosidic bonds between the sugar moieties, (ii) amidases cleave the bond between MurNAc and the stem peptide, and (iii) endopeptidases, which target peptide bonds within the stem peptides or interpeptide bridges (1, 10).

Many streptococci have the ability to take up extracellular DNA and incorporate it into their genome through homologous recombination by entering a physiological state known as natural competence for genetic transformation (11–16). This is important for genetic plasticity and evolution, which is believed to increase their capacity to handle environmental challenges (17–20). Competence in streptococci is a tightly regulated process, often induced by environmental cues, and is primarily controlled by quorum-sensing mechanisms involving the ComCDE (mitis and anginosus group streptococci) or the ComRS (pyogenic, salivarius, mutans and bovis group streptococci) regulatory systems. These pathways involve the competence-inducing peptides CSP and XIP, respectively. The competence-inducing peptides activate signaling cascades that lead to the expression of early and late competence genes, which are essential for the uptake and integration of extracellular DNA into the genome (11).

The process of natural transformation is best studied in *S. pneumoniae* (reviewed in (reviewed in 20, 21-23). During competence, *S. pneumoniae* expresses a lytic murein hydrolase called CbpD, which is utilized by competent cells to specifically kill and lyse non-competent pneumococci and closely related bacteria, resulting in the release of their cytoplasmic contents; a predatory mechanism known as fratricide (3, 24). Fratricide is proposed to supply the competent cells with homologous DNA that is likely to integrate successfully into their genome (3, 20, 24, 25). CbpD consists of three domains: an N-terminal cysteine, histidine-dependent amidohydrolase/peptidase (CHAP) domain, which acts as an amidase or endopeptidase, two peptidoglycan-binding SH3b domains, and a C-terminal choline binding domain (CBD) that binds to choline residues on teichoic acids within the cell wall (24, 26). Beyond *S. pneumoniae*, putative fratricins have been identified in other streptococcal species based on their genomic positions and promoters (25). Some species possess CbpD-like fratricins, such as those in the mitis group streptococci (25), while others have putative CHAP-domain-containing fratricins that either lack the CBD, such as CrfP in *Streptococcus suis* (27), or contain alternative cell wall binding domains. Examples of the latter include LytF in *S. mutans* and *Streptococcus sanguinis*, which harbor so-called Bsp-domains (28, 29). Additionally, putative fratricins in the pyogenic group (e.g., *S. dysgalactiae, S. pyogenes*) and salivarius group streptococci (e.g., *Streptococcus thermophilus, Streptococcus vestibularis*) possess a fully uncharacterized C-terminal domain (25). Some streptococci also encode non-CHAP domain-containing putative fratricins such Zoocin A in *Streptococcus agalactiae* and *Streptococcus equi,* which feature a distinct, uncharacterized cell binding domain combined with an M23-like catalytic domain shown to function as an endopeptidase (25, 30–32).

As mentioned above, bacteria require a dedicated immunity mechanism to protect themselves from committing suicide during expression of the competence-induced murein hydrolases. In competent *S. pneumoniae*, self-protection is mediated through expression of the immunity protein ComM, a multi-transmembrane spanning protein closely related to the CPBP-family of intramembrane proteases (33). ComM has been shown to delay cell division; however, the precise mechanism by which ComM confers immunity in *S. pneumoniae* remains undetermined (34–36). Additionally, for Zoocin A, a Fem-transferase-like protein has been implicated in immunity by modifying the length of the interpeptide bridge (37, 38).

The pyogenic group of streptococci includes important pathogens such as *S. pyogenes* and *S. dysgalactiae*. *S. pyogenes* is responsible for a range of human infections, from mild conditions like pharyngitis to severe diseases such as septicemia, streptococcal toxic shock syndrome (STSS), and necrotizing fasciitis (39), while *S. dysgalactiae* causes mastitis in cows as well as various infections in humans, including superficial skin infections, STSS, meningitis, and endocarditis (40). The role of competence and natural transformation in these bacteria remains obscure. In a recent study, we found that the putative fratricin encoding gene of *S. dysgalactiae* was indeed upregulated during competence (41). This enzyme, comprising a CHAP domain, an SH3b domain, and a C-terminal conserved domain (CCD) of unknown function (Fig. 1A), has not previously been studied.

**Fig. 1.**
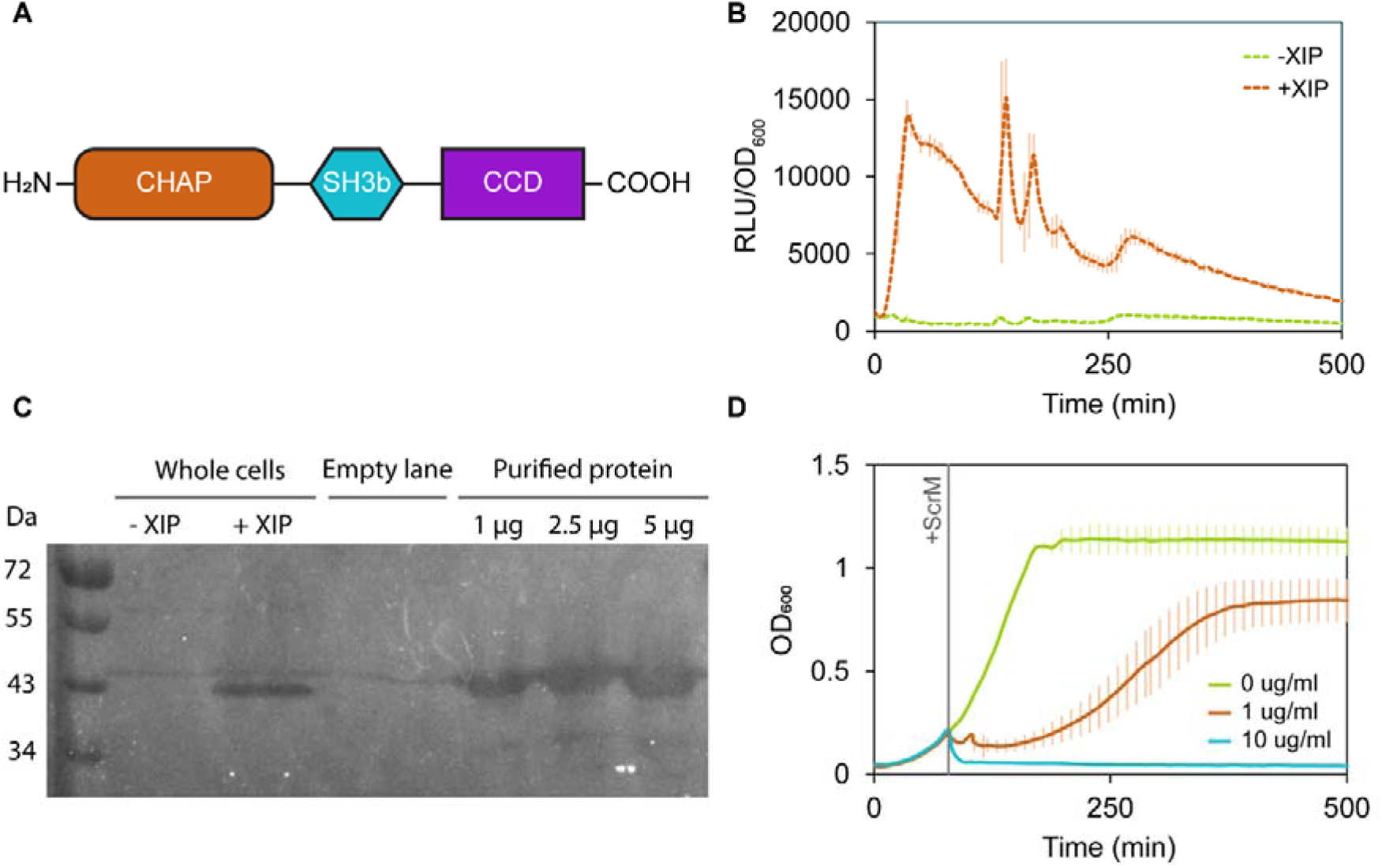
Characterization of ScrM activity. **(A)** Schematic representation of the domain organization of ScrM. Mature ScrM contains an N-terminal CHAP domain, an SH3b domain, and an uncharacterized conserved C-terminal domain (CCD) of the lytic exoenzyme target recognition domain superfamily (IPR038263). **(B)** Activation of P*_scrM_* during competence. Luminescence was monitored in *S. dysgalactiae* cells harboring a transcriptional reporter (firefly luciferase) under the control of the P*_scrM_* promoter (strain MM420). Cells were either exposed to the competence-inducing peptide XIP (orange line) or left untreated (green line). Luminescence values were normalized to cell density (OD_600_) and are presented as RLU/OD_600_. Data represent the mean and standard deviation of three technical replicates, and are representative of three independent experiments, demonstrating a rapid induction of the P*_scrM_* promoter upon competence induction. **(C)** Zymogram analysis of ScrM murein hydrolase activity. The murein hydrolase activity was assessed in whole cell extracts from *S. dysgalactiae* and for purified ScrM. Heat-inactivated *S. dysgalactiae* cells were incorporated into the SDS-PAGE separation gel, and lytic activity was observed as clear bands within the opaque gel. Competence-induced cells were harvested 1-hour post-XIP addition (+XIP), and non-induced cells were analyzed in parallel (-XIP). Purified ScrM (1 µg, 2.5 µg, 5 µg) was also included in the zymogram. The zymogram was developed over 2 hours. **(D)** The lytic effect of purified ScrM on exponentially growing *S. dysgalactiae* cells was quantified by measuring OD_600_ at 10-minute intervals. ScrM (0 µg/ml, 1 µg/ml or 10 µg/ml) was added to cultures at an OD_600_ of ∼ 0.2. The data represent the mean of three technical replicates, with error bars indicating the standard deviation. The data are representative of three independent experiments.

In this work, we investigated the lytic function of this enzyme, named ScrM (for <u>S</u>treptococcal <u>C</u>ompetence <u>R</u>egulated <u>M</u>urein hydrolase), its cell surface binding properties and the mechanism underlying self-immunity in *S. dysgalactiae*. We found that ScrM exhibits lytic activity against streptococci of the pyogenic group (*S. dysgalactiae, S. pyogenes, S. agalactiae*) and the salivarius group (*S. thermophilus, S. vestibularis*). Additionally, our study revealed that a competence-induced Fem-transferase-like protein modifies the amino acid composition of the interpeptide bridge in the PG, providing self-protection against ScrM.

## Results

### *S. dysgalactiae* encodes a competence induced murein hydrolase

In our previous work, we identified a CHAP-domain-encoding gene (Text S1) that was upregulated in response to induction with the competence-stimulating peptide XIP in RNA-seq experiments (41). This protein, which contains an uncharacterized conserved C-terminal domain (CCD) in addition to an N-terminal signal sequence, a CHAP domain and an SH3b domain (Fig. 1A), has not previously been studied. However, it has been hypothesized to function as a competence-induced fratricin in *S. dysgalactiae* and other pyogenic group streptococci (25). In this study, we aimed to characterize this putative hydrolase, henceforth designated ScrM. To confirm that ScrM is indeed competence regulated, we placed a transcriptional reporter (*luc* encoding firefly luciferase) under the control of the *scrM* promoter (P*_scrM_*). Growth (OD_600_) and luminescence were monitored over time (Fig. 1B) in the presence or absence of XIP. A rapid increase in luminescence was observed approximately 10 minutes after addition of XIP, confirming that the P*_scrM_*promoter is activated during competence.

Given the presence of the CHAP domain, which is associated with murein hydrolase activity, we hypothesized that ScrM functions in the degradation of peptidoglycan. To test this, we assessed the murein hydrolase activity of competence-induced cells using zymography using heat-killed cells of non-competent *S. dysgalactiae* as ScrM-substrate. A clear band corresponding to the molecular mass of the mature ScrM protein was observed in the gel (Fig. 1C), showing that ScrM is a novel competence-dependent murein hydrolase.

The *scrM* gene was subsequently cloned and overexpressed in *Escherichia coli* (the predicted N-terminal signal peptide of ScrM was replaced with a 6x His-tag), followed by purification using Immobilized-Metal Affinity Chromatography (IMAC). All experiments involving purified ScrM were performed using the His-tagged version. The purified protein had the expected molecular mass of 39.9 kDa as determined by SDS-PAGE (Fig. S1) and displayed muralytic activity in zymogram analysis (Fig. 1C) as well as in liquid culture assays (Fig. 1D). More lysis was observed with higher concentrations of ScrM, demonstrating a dose-dependent effect.

### ScrM is active against pyogenic and salivarius group streptococci

To test the target range of ScrM, we treated 11 different streptococcal species with purified ScrM in liquid cultures, including members of the pyogenic, mitis, suis, bovis, salivarius, and mutans streptococcal subgroups (Table S1). The species were grown in microtiter plates to an OD_600_ of ∼0.1-0.2, after which ScrM was added to final concentrations of 1 µg/ml and 10 µg/ml. Due to differences in growth medium preferences, some species were grown in C-medium, while others were grown in BHI. Our results, presented as relative change in optical density after ScrM treatment, demonstrated that ScrM exhibits significant lytic activity specifically for species within the pyogenic (*S. dysgalactiae*, *S. pyogenes* and *S. agalactiae*) and salivarius (*S. thermophilus* and *S. vestibularis*) groups (Fig. 2). Additionally, we observed that ScrM was more potent in C-medium compared to BHI when examining its activity against *S. dysgalactiae* suggesting that the lytic activity of ScrM is influenced by the growth medium used.

**Fig. 2.**
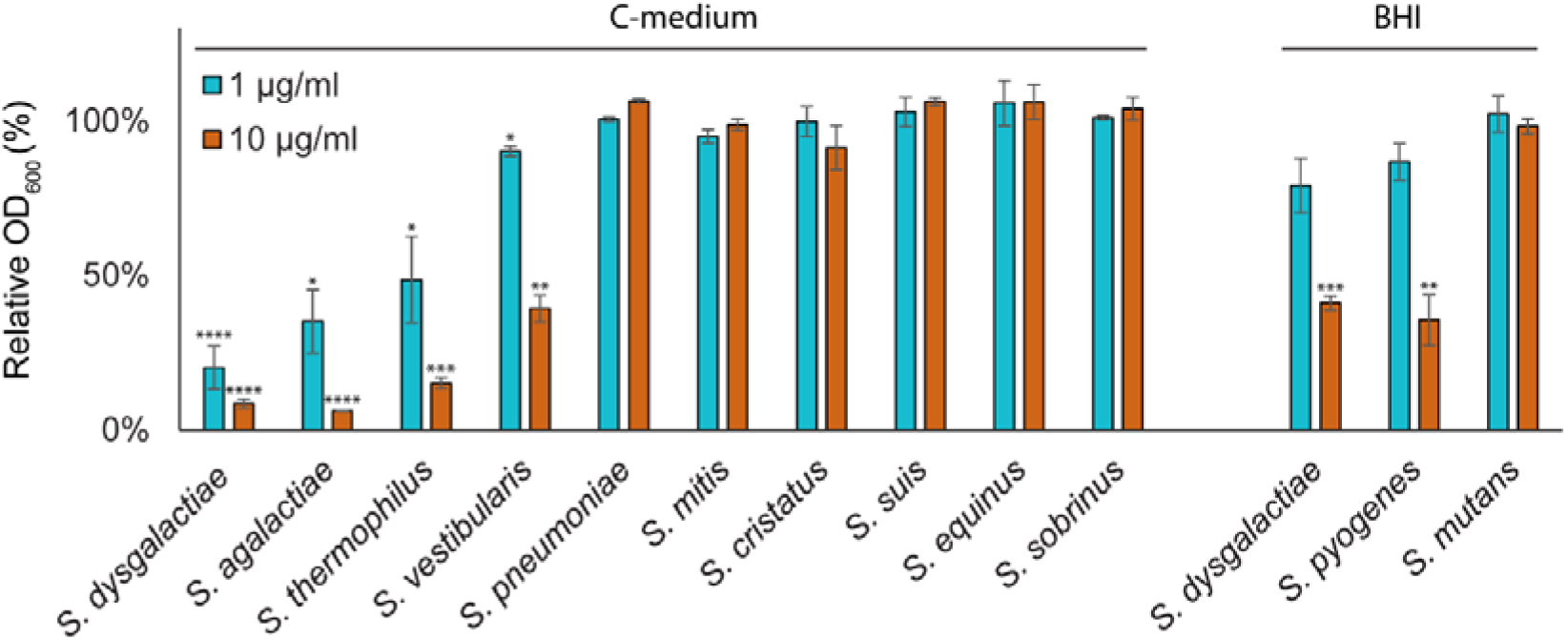
Target range of ScrM against streptococci. The effect of ScrM on growth of various streptococcal species was assessed by measuring the relative decrease in OD_600_ of cultures treated with ScrM. Strains were grown to an OD_600_ of ∼ 0.1 – 0.2 in either C-medium or BHI before ScrM was added to final concentrations of 1 µg/ml (blue bars) or 10 µg/ml (orange bars). The y-axis represents the percentage of OD_600_ compared to the untreated control, reflecting the extent of cell lysis. Columns represent the mean of at least three independent experiments, with error bars indicating standard deviation. Statistical significance relative to the untreated control was calculated using a one-sample t-test and is indicated by asterisks: * *p* < 0.05, ** *p* < 0.01, *** *p* < 0.001, **** *p* < 0.0001.

### ScrM targets the division zone

In addition to the catalytic CHAP domain, ScrM has an SH3b domain and a CCD domain. The CCD domain remains uncharacterized, whereas SH3b domains are described as cell-wall binding, often found in proteins with peptidoglycan hydrolase activity (26, 42, 43). Target cell recognition by ScrM is most probably mediated by the SH3b and CCD domains. To examine the cell surface binding specificity of ScrM, we therefore created an sfGFP-SH3b-CCD fusion protein (CHAP replaced with sfGFP) that was overexpressed and purified similar to ScrM described above (Fig. S1). Fluorescence microscopy of non-competent *S. dysgalactiae* cells revealed that the fusion protein predominantly targeted the division zone of the cells, with additional binding observed along the remaining cell surface, as evidenced by heat map analysis (Fig. 3A). The binding pattern was consistent across other ScrM-sensitive species, such as *S. pyogenes* (Fig. S2). However, no localized binding was detected in the non-sensitive *S. pneumoniae*, and the overall binding to pneumococcal cells was low (Fig. S2). Notably, the binding of sfGFP-SH3b-CCD varied between *S. dysgalactiae* cells of different stages of cell division (Fig. 3C); weak binding of the fusion protein was observed around the entire cell before cell division was initiated. For cells with early septum formation, sfGFP-SH3b-CCD binding was primarily detected at the division zone. Upon completion of cell division and the splitting of daughter cells, the sfGFP-SH3b-CCD fusion protein was found to bind in the area between the newly divided cells (Fig. 3C).

**Fig. 3.**
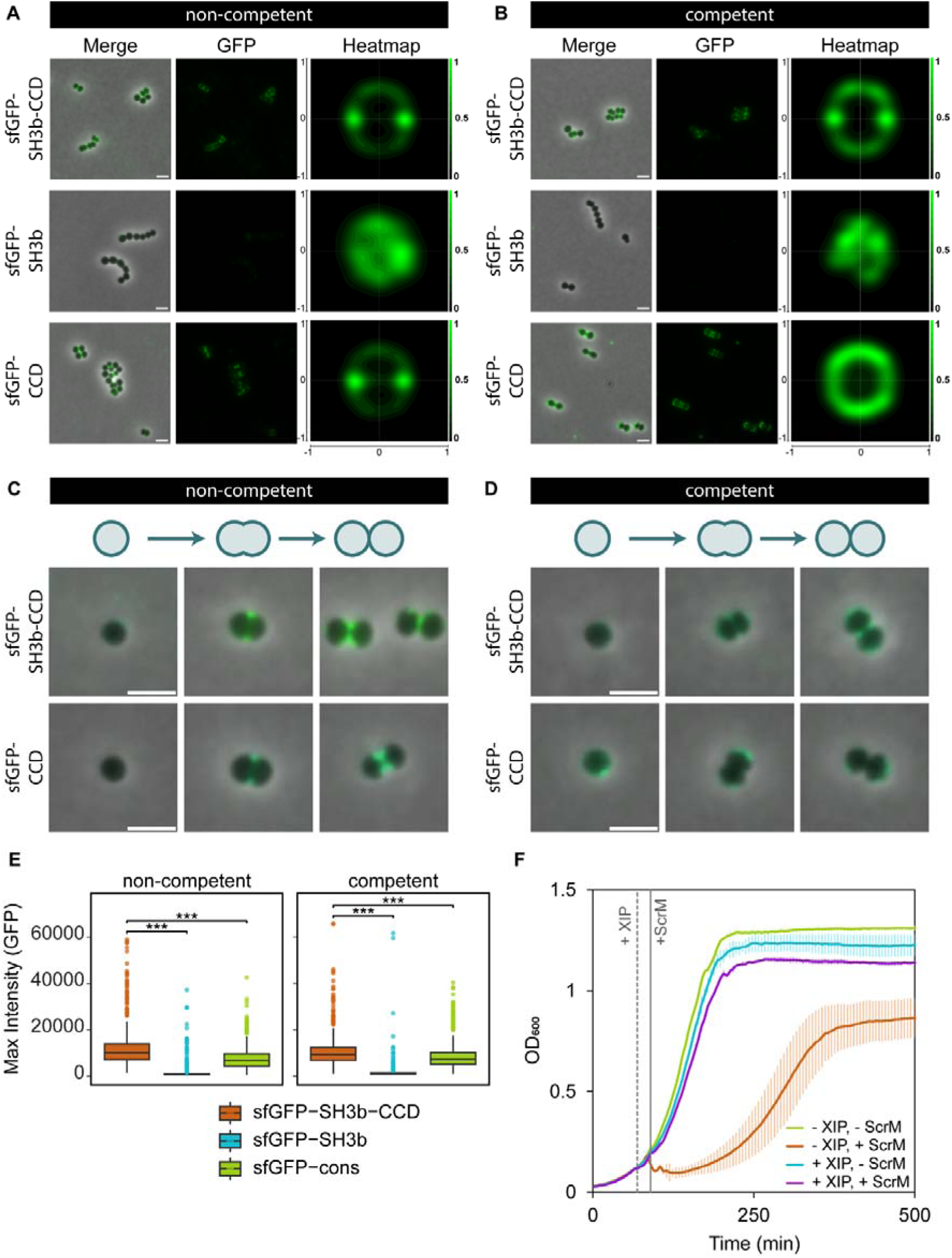
Localization and activity of ScrM in competent and non-competent cells. Localization of sfGFP-SH3b-CCD, sfGFP-SH3b, and sfGFP-CCD fusion proteins on **(A)** non-competent and **(B)** competent *S. dysgalactiae* cells. The number of cells used to generate heatmaps for non-competent cells are sfGFP-SH3b-CCD (n = 1086), sfGFP-SH3b (n = 1521), sfGFP-CCD (n = 1127), and for competent cells sfGFP-SH3b-CCD (n = 1135), sfGFP-SH3b (n = 1015), and sfGFP-CCD (n = 1019). Scale bars represent 2 µm. **(C)** Microscopy images showing the localization of sfGFP-SH3b-CCD and sfGFP-CCD on non-competent *S. dysgalactiae* cells captured at different stages of cell division. The top panel depicts different cell cycle phases, corresponding to the stages of cell division observed in the images. Scale bars represent 2 µm. **(D)** Similar as panel C but with competent cells. **(E)** Quantification of GFP-signal intensity of non-competent and competent *S. dysgalactiae* cells labelled with sfGFP-SH3b-CCD, sfGFP-SH3b, or sfGFP-CCD fusion proteins. Statistical significance was determined using a Kurskal-Wallis test, followed by Dunn’s post-hoc test. Significance levels are indicated as follows: * *p_adj_* < 0.05, ** *p_adj_* < 0.01, *** *p_adj_* < 0.001. **(F)** Competence-induced immunity to ScrM in *S. dysgalactiae*. The growth of *S. dysgalactiae* was monitored for competent (+XIP) and non-competent (-XIP) cells, with and without the addition of purified ScrM (1 µg/ml). OD_600_ was measured at 10-min intervals. XIP was added at OD_600_ ∼ 0.1 to induce competence, followed by the addition of ScrM 20 minutes post-XIP induction. Data represent the mean of three technical replicates, with error bars indicating standard deviation.

Next, we investigated the individual contribution of the SH3b and the CCD domains to the septal binding by purifying sfGFP-SH3b and sfGFP-CCD (Fig. S1). Labeling cells with these sfGFP-fusions demonstrated that only a small fraction of cells exhibited binding of sfGFP-SH3b (5.7%, n = 87/1521) compared to sfGFP-SH3b-CCD (54.8%, n = 595/1086) and sfGFP-CCD (50.2%, n = 566/1127). Moreover, the sfGFP-SH3b fusion protein exhibited a diffuse binding pattern without clear localization to the division zone and the maximum GFP intensity was considerably lower than that observed with sfGFP-SH3b-CCD and sfGFP-CCD (Fig. 3A, Fig. 3E). In contrast, the sfGFP-CCD fusion targeted the division zone similar to sfGFP-SH3b-CCD (Fig. 3A and C) but with a somewhat reduction in overall GFP signal on cells (Fig. 3E). These findings suggest that the targeting of ScrM to the division zone is predominantly dependent on the CCD domain, while the SH3b domain may play a supportive role in enhancing binding.

The CCD domain has not previously been characterized but is conserved among putative competence induced murein hydrolases in the pyogenic and salivarius group (25). Multiple sequence alignments of CCD domains across nine streptococcal species revealed a conserved four-amino acid motif, LAGG (Fig. S3A). Based on its surface positioning in the predicted ScrM 3D-structure (Fig. S3B, Fig. S3C), we hypothesized that the LAGG motif is important for mediating binding to the target cell or division zone. To test this, we purified a mutant sfGFP-CCD^mut^ fusion protein by substituting the glycines of the LAGG motif into alanines (Fig. S1). Localization analysis showed that the mutant protein exhibited weak, diffuse binding across the cell surface, with no specific localization to the division zone (Fig. S3D). Additionally, introducing the same substitutions in the full-length ScrM (ScrM^G302A,G303A^) inactivated the lytic activity in liquid cultures, comparable to the iodoacetamide-inactivated ScrM control (Fig. S3E). These findings show that the lytic function of ScrM depends on binding to the division zone of target cells which is determined by the CCD domain and particularly the LAGG motif.

### Competence results in self-immunity to ScrM

Given the notable lytic activity of ScrM towards *S. dysgalactiae*, competent cells expressing ScrM must possess a mechanism to prevent self-killing. To investigate whether *S. dysgalactiae* exhibits competence-induced immunity to ScrM, we compared lysis of non-competent and competent cells after addition of purified ScrM (Fig. 3F). Our results indeed demonstrate that competent cells, which were subjected to XIP for ∼20 minutes, are immune to ScrM, whereas non-competent cells remain sensitive.

To explore the mechanism underlying this immunity, we analyzed the localization of the GFP-fused full-length protein (sfGFP-SH3b-CCD), as well as the individual cell wall-binding domains (sfGFP-SH3b and sfGFP-CCD) on competent cells of *S. dysgalactiae*. The sfGFP-SH3b-CCD binding to competent cells (Fig. 3B) was clearly different from the non-competent cells (Fig. 3A); while still displaying enriched binding at the division zone, a larger fraction of the protein bound to the cell periphery. Even more dramatically, the sfGFP-CCD fusion protein fully lost its distinct localization to the division zone of competent cells. Heatmap analysis indicated that the binding was, in fact, reduced at division zones compared to the cell periphery (Fig. 3B and 3D). As expected, the sfGFP-SH3b fusion protein displayed similar binding pattern on competent cells and on non-competent cells, with weak and nonspecific binding (Fig. 3B, Fig. 3E).

### ScrI mediates competence-dependent immunity to ScrM

Since ScrM is expressed during competence and cell immunity is competence-dependent, it is likely that the corresponding immunity mechanism is activated early during competence to prevent self-lysis. Our previous RNA-seq analysis (41) identified several genes of unknown function that are upregulated during competence. We reasoned that since the lytic function of ScrM relies on its specific binding to the division zone of target cells, the associated immunity mechanism might be linked to cell wall synthesis or its modulation. No homolog to the *S. pneumoniae* CbpD-immunity protein ComM (33–35) was found among the upregulated genes in *S. dysgalactiae*. We did, however, identify one gene with high sequence similarity to genes encoding Fem transferases. The gene, which we have named *scrI* (<u>S</u>treptococcal <u>C</u>ompetence-<u>R</u>egulated Immunity factor, Text S1), harbors a CIN-box in the promoter region, typically associated with competence-induction (41) (Fig. 4A). Although *scrI* is expressed at low levels under non-competence conditions, the gene becomes highly upregulated during competence (41).

**Fig. 4.**
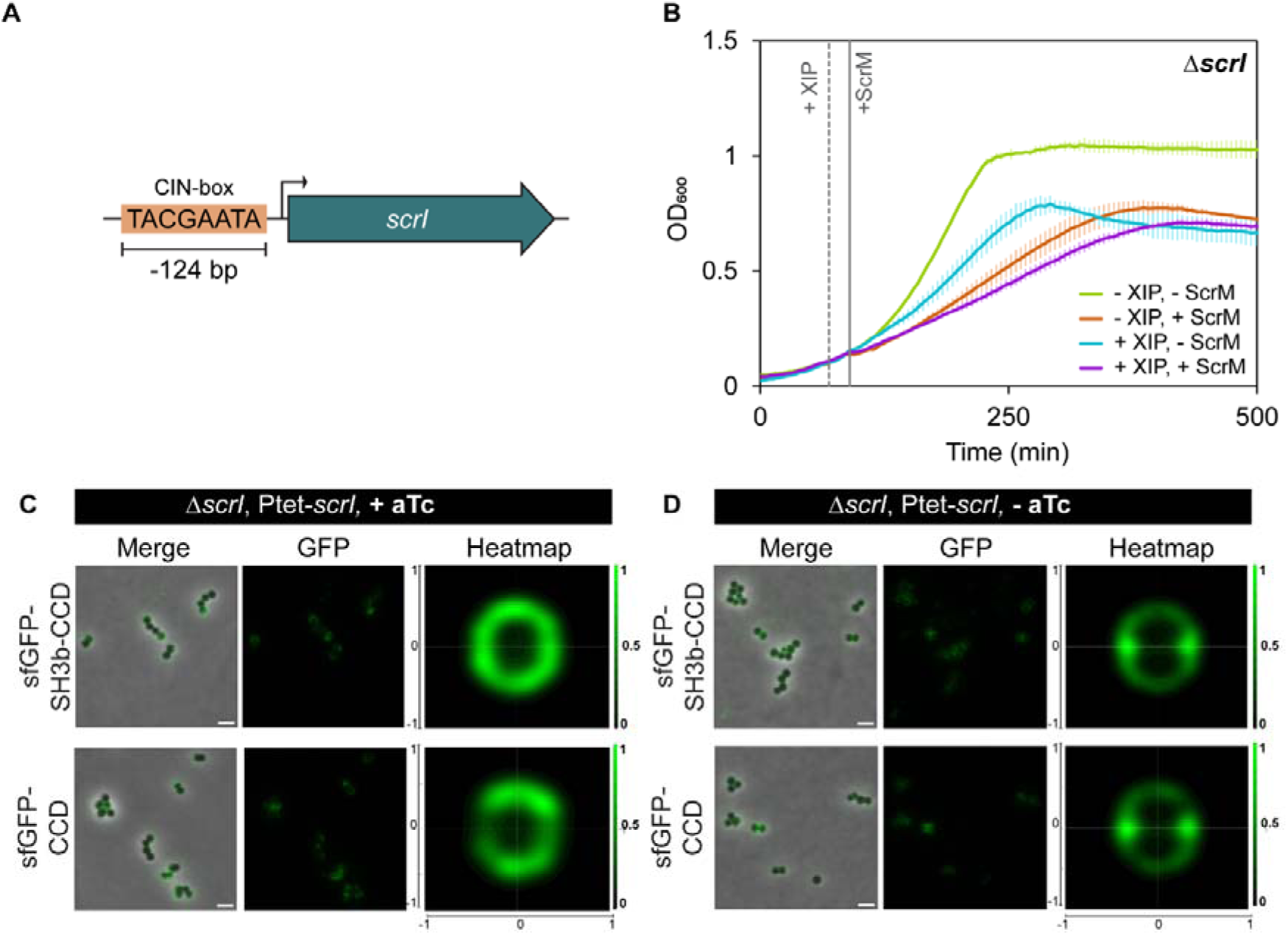
ScrI confers immunity to ScrM. **(A)** Schematic diagram of the *scrI* gene with the CIN-box sequence 124 bp upstream of the coding sequence **(B)** Growth curves of non-competent and competent (250 ng/ml XIP) Δ*scrI* cells (strain MM463) exposed to 0 or 1 µg/ml purified ScrM. XIP was added at an OD_600_ of ∼ 0.1 and ScrM was added 20 min post XIP-induction. OD_600_ was measured at 10-minute intervals. Data represent the mean of three technical replicates, with error bars indicating standard deviation, and is representative of three independent experiments. **(C)** Microscopy analysis showing the binding pattern of sfGFP-SH3b-CCD and sfGFP-CCD fusion proteins in the Δ*scrI* mutant with ectopic expression of ScrI (strain MM481, Δ*scrI::kan*, pFD116-P_tet_-*scrI*). Ectopic expression of ScrI was induced by adding a final concentration of 100 ng/ml aTc. Heatmap analysis show the binding patterns of these fusion proteins based on n = 1067 and n = 1034 cells, respectively. Scale bars represent 2 µm. **(D)** Binding pattern of sfGFP-SH3b-CCD and sfGFP-CCD fusion on the Δ*scrI* mutant (strain MM481, Δ*scrI::kan*, pFD116-P_tet_-*scrI*) without induction of ectopic ScrI expression,. Heatmap analysis indicate the binding patterns of these fusion proteins based on n = 1414 and n = 1114 cells, respectively. Scale bars represent 2 µm.

To investigate whether ScrI is involved in immunity to ScrM, we created a *scrI* knockout mutant (Δ*scrI::kan*) and subjected both competent and non-competent cells to purified ScrM (Fig. 4B). Strikingly, in the absence of ScrI, competence induction did not confer any immunity, as evidenced by the strong inhibition of growth when the mutant was exposed to ScrM (Fig. 4B). Additionally, when competence was induced in the Δ*scrI* mutant, there was a notable reduction in growth compared to the non-induced strain, even without added ScrM. This reduction is likely due to the lack of self-immunity to the ScrM expressed during competence, resulting in self-killing. The phenotype could be complemented by tetracycline inducible expression of ScrI from a plasmid (Fig. S4). Together, these results demonstrate that ScrI is indeed essential for effective immunity against ScrM during competence. It should be noted that the Δ*scrI* mutant exhibited reduced growth compared to the wild type, even under non-competent conditions, suggesting that low levels of ScrI expression is needed for normal growth.

### ScrI alters ScrM binding pattern

Previous observations indicated that ScrM does not bind to non-susceptible cells (Fig. S2), suggesting that immunity may be linked to the binding specificity of ScrM. We hypothesized that ScrI might influence the surface of competent cells, potentially playing a role in mediating immunity by altering ScrM’s interaction with the cell surface.

To test this, we performed microscopy analyses where ScrI was ectopically expressed in the Δ*scrI* mutant. We then observed distinct changes in the binding patterns of the sfGFP-fusion proteins (Fig. 4C); sfGFP-SH3b-CCD, which typically binds to the division zone on sensitive cells, lost this specificity and was instead distributed evenly across the entire cell surface. Moreover, sfGFP-CCD exhibited an even more striking alteration in binding pattern, i.e. almost entirely being excluded from the division zone. In contrast, the Δ*scrI* mutant without induction of ectopic expression of ScrI showed similar binding patterns to those observed in the non-competent wild type with binding predominantly at the division zone (Fig. 4D). Together, these findings support the role of ScrI in modulating the cell envelope of competent cells in such a way that ScrM no longer can bind properly to the cell surface, which is essential for its lytic activity.

### ScrI protects against ScrM by modulating the peptidoglycan interpeptide bridges

To investigate whether ScrI induced modifications in the cell wall, we performed LC-MS/MS to analyze the muropeptide composition of peptidoglycan derived from competent ScrI-deficient cells (non-immune) and the Δ*scrI* mutant ectopically expressing ScrI (immune). Comparison of the LC chromatograms revealed distinct differences in the presence and intensity of muropeptide peaks between the two strains. Some muropeptides were found to be present only in the strain ectopically expressing ScrI. The ScrI-deficient cells exhibited peaks corresponding to specific muropeptide masses (e.g., *m/z* = 584, *z* = 2), while ScrI-proficient cells displayed additional peaks having a 30 Da increase in mass (e.g., *m/z* = 599, *z* =2) along with a 10-fold reduction in the original peak intensities (Fig. 5A). A similar comparison of the muropeptide pair of *m/z* = 555/570 (*z* = 2) can be found in Fig. S5A. These observations suggest that ScrI induces modifications in the peptidoglycan, specifically amino acid substitutions, corresponding to a 30 Da mass shift.

**Fig. 5.**
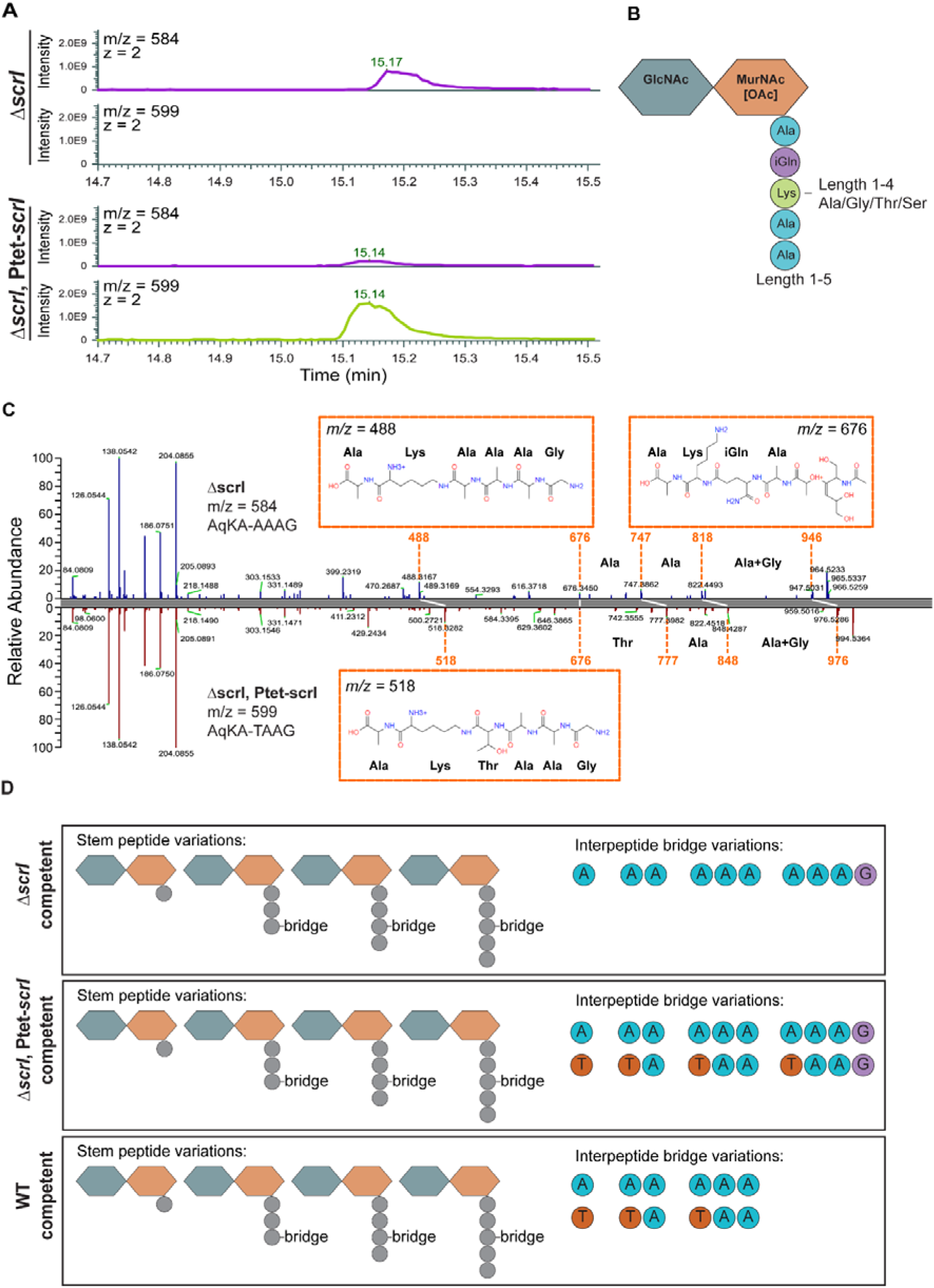
LC-MS/MS analysis of peptidoglycan. **(A**) LC-MS/MS chromatograms showing the comparison of muropeptide peaks in competent cells of the Δ*scrI* mutant (top, strain MM463) and the Δ*scrI* mutant ectopically expressing ScrI (bottom, strain MM481). The Δ*scrI* mutant displays a peak at *m/z* = 584 (*z* = 2), while the strain expressing ScrI shows a 10-fold reduction of this peak but instead an additional peak at *m/z* = 599 (*z* =2), indicating a total 30 Da increase in mass. Notably, this peak is not present in the Δ*scrI* mutant **(B)** Schematic representation of the parameters used for generating the *in silico* library with PGN_MS2 (62). GlcNAc: N-Acatylglucosamine; MurNAc: N-Acetylmuramic acid, [OAc]: O-Acetylation; Ala: Alanine; iGln: iso-γ-Glutamine; Lys: Lysine. **(C)** LC-MS/MS spectra for *de novo* analysis of mass peaks *m/z* = 584 in the competent Δ*scrI* mutant (top, strain MM463) highlighting signature ions representing a fragment of the stem peptide without a bridge (*m/z* = 676) and the same fragment with increasing bridge components (*m/z* = 747, 818, 946), verifying the AAAG bridge. A second signature ion is the *m/z* = 488, which represent a stem-fragment with AK plus the entire bridge. In the competent ScrI expression strain (bottom, strain MM481) we see the same stem fragment without bridge (*m/z* = 676), but a shifted series of +30 Da compared to the Δ*scrI* mutant, and it is clear that the + 30 Da shift occurs in position one indicating an Ala➔Thr modification. Similarly, the stem-fragment with AK plus the entire bridge is now 30 Da heavier. *m/z* = 584 was determined to be AqKA-[3-NH2-AAAG], while *m/z* = 599 was determined to be (AqKA-[3-NH2-TAAG]. A: Alanine, q: γ-iso-Glutamine; K: Lysine; G: Glycine. **(D)** Proposed models of peptidoglycan structures for the competent Δ*scrI* mutant, the ScrI expression strain and the wildtype (WT). A: Alanine; T: Threonine; G: Glycine.

To identify the nature of these 30 Da differences, we compared the experimental MS/MS spectra with an *in silico* generated MS/MS library of predicted muropeptides and corresponding MS/MS spectra (see Methods for details). The *in-silico* predictions were based on the observed 30 Da mass differences, as well as known variation in interpeptide bridges reported for *S. dysgalactiae* and related species(7) (Table S1), with the input parameters for the library generation depicted in Fig. 5B. Comparison of the experimental and predicted MS/MS spectra revealed no major differences in stem peptide composition or glycan chain modifications between the strains (Table S2, Table S3). All strains predominantly displayed muropeptides with tripeptides, tetrapeptides and pentapeptides in the stem, with occasional O-acetylation on MurNAc (Table S2, Table S3).

However, notable differences were observed in the interpeptide bridges between the Δ*scrI* and the ScrI expression strains (Table S2, Table S3). In the Δ*scrI* mutant, interpeptide bridges were primarily composed of Ala residues, such as [Ala], [Ala-Ala] and [Ala-Ala-Ala], with some interpeptide bridges containing Gly at the fourth position ([Ala-Ala-Ala-Gly]). A subset of these muropeptides, including [Ala-Ala-Ala-Gly], was manually inspected, further confirming this organization (Fig. 5C). In contrast, the ScrI expression strain exhibited a broader variety of interpeptide bridges, most notably including interpeptide bridges with Thr located in the first position, such as [Thr-Ala-Ala-Gly] and [Thr-Ala-Ala] (Fig. 5C, Fig. S5B).

We extended this analysis to wild type cells induced to competence, which, similar to the ScrI expression strain, showed a combination of interpeptide bridges with both Ala and Thr in the first position (Table S4), including [Ala-Ala-Ala] and [Thr-Ala-Ala]. We did, however, not detect interpeptide bridges with Gly in the fourth position in the WT. Taken together, these results suggests that ScrI deficient cells (non-immune) harbors peptidoglycan consisting of interpeptide bridges without Thr, while during competence (immune cells) a proportion of the interpeptide bridges are modified to contain Thr in the first position (Fig. 5D). This modification likely disrupts the ability of ScrM to bind effectively the division zone, ultimately protecting the cells from ScrM-mediated lysis.

## Discussion

In this study, we characterized the competence-induced murein hydrolase ScrM in *S. dysgalactiae* and elucidated the mechanism underlying self-immunity conferred by the Fem-transferase-like protein ScrI. ScrM displayed specific muralytic activity against streptococci belonging to the pyogenic and salivarius subgroups (Fig. 2), aligning well with the idea that ScrM functions as a fratricin for pyogenic streptococci.

ScrM specifically binds the division zone of susceptible cells, and the uncharacterized CCD domain is critical for this binding (Fig. 3A and 3C). The CCD domain likely recognizes specific structures that are enriched, more accessible or only present in newly synthesized cell wall. The SH3b domain of ScrM displayed relatively poor binding to cells compared to the CCD domain and did not localize to the division zone (Fig. 3A). This binding pattern is similar to SH3b domains of other streptococcal murein hydrolases, like CbpD in *S. pneumoniae* and CrfP *S. suis*, which bind unknown targets in PG (26, 27). The SH3b domain of lysostaphin in *S. aureus*, on the other hand, have been shown to specifically target the pentaglycine interpeptide bridges in PG and structural data on SH3b domains of phage lysins also suggest binding to peptide side chains in PG (42–45). One might therefore propose that the SH3b domain of ScrM, while not sufficient alone for proper binding to cells (Fig. 3A), has a similar binding preference, which contributed to slightly enhanced binding of sfGFP-SH3b-CCD to target cells (Fig. 3E).

It should be noted that some murein hydrolases also have domains that bind cell wall-associated polysaccharides instead of PG, such as the choline-binding domains of CbpD in *S. pneumoniae*, which mediates specific binding to choline on wall teichoic acids (24, 26). Many streptococci, including *S. dysgalactiae* and other ScrM-sensitive species, harbor cell wall-associated rhamnose-glucose polysaccharides (RGPs) instead of wall teichoic acids (46) (Table S1). However, these are also found among non-sensitive species, such as *S. mutans* (46), suggesting they are unlikely to be the primary binding targets for the CCD domain of ScrM. In this context, it is also interesting to note that wall teichoic acids have been shown to regulate the binding of PG hydrolases by acting as steric hindrances. For example, autolysins like Atl in *S. aureus* localizes to the division zone because binding to peptidoglycan elsewhere is inhibited due to the presence of teichoic acids (47). In *S. dysgalactiae*, RGPs could potentially function similarly, however, the subcellular localization of these polysaccharides in streptococci remains to our knowledge largely unexplored. Our data thus instead support a model where CCD primarily recognize structures in PG.

We demonstrated that competence-induced cells exhibit immunity to ScrM by expressing the Fem-transferase-like immunity factor ScrI, previously found to be upregulated during competence (41). ScrI shares similarities to the lipid II peptidyl transferases, such as MurM in *S. pneumoniae,* which catalyze the addition of the first amino acid in the interpeptide bridge (48, 49). Our LC-MS data suggest that while Ala is typically incorporated as the first amino acid in the interpeptide bridge in *S. dysgalactiae*, likely by the constitutively expressed FemX (Stdys021_00117), ScrI appears to incorporate Thr in place of Ala at this position (Fig. 5C, Fig. 5D, Fig. S5A, Fig. S5B). The increased presence of Thr in the interpeptide bridges is accompanied by reduced septal binding of CDD and SH3-CDD (Fig. 3B, Fig. 3D), suggesting that the CCD binds preferentially to PG containing Ala rather than Thr. Interestingly, the loss of septal binding is even more pronounced when ScrI is ectopically expressed compared to its natural expression levels during competence (Fig. 4C). The LC-MS data demonstrate that there is a higher proportion of Thr-containing muropeptides in these cells (Table S3, Table S4), showing that increased levels of Thr in interpeptide bridges indeed correlates with reduced binding of ScrM (Fig 4C). Exactly how replacement of Ala with Thr can compromise ScrM-binding needs to be further examined by structural studies. Interestingly, however, resistance to murein hydrolases from Fem-transferase mediated alterations of interpeptide bridges has been suggested in different species previously (50–54). For example, the FemABX-like protein Zif in *S. equi* confers immunity to the murein hydrolase Zoocin A through an additional Ala in the interpeptide bridge, and the results suggested that this impacts both binding and hydrolytic activity (37). In *S. aureus*, it has been shown that FmhA and FmhC incorporate Ser residues into the interpeptide bridges to confer immunity to lysostaphin (54). Together with the results presented here, this shows that expression of dedicated Fem-transferases to alter interpeptide bridge composition is a mechanism used to for protection against specific murein hydrolases across different species and genera.

The muralytic activity of ScrM is mediated by the well-known CHAP domain. These domains can function as amidases, cleaving the amide bond between the MurNAc and the primary Ala in the pentapeptide, or as an endopeptidase, cleaving peptide bonds between amino acid residues in the peptide bridges (55). The exact bond(s) targeted by ScrM remains to be fully elucidated but is likely influenced by the composition of the PG. While our data support a model where immunity is mediated by reduced binding of ScrM to the division zone due to Thr-containing interpeptide bridges, it is also possible that incorporation of Thr residues creates a PG that is a less optimal substrate for the CHAP domain, rendering the cells immune to lysis. Interestingly, ScrM still binds to the remaining cell surface in immune cells. This suggest that the lytic function of ScrM may be inhibited outside the division zone or that its enzymatic target is newly synthesized PG in the septal region. The latter is similar to what has been reported for the pneumococcal fratricin CbpD (26, 56). Furthermore, structural modifications, such as O-acetylation of MurNAc, detected in both ScrI-expressing and ScrI-lacking cells (Table S2), or the presence of RGPs as mentioned above, could play a protective role by reducing the susceptibility of PG to murein hydrolases. O-acetylation of MurNAc is known to play an important role for resistance to lysozyme in many bacteria, as well as to influence the activity of bacterial autolysins (57, 58).

Here we have demonstrated that competent *S. dysgalactiae* expresses a murein hydrolase, ScrM, capable of lysing close relatives, in addition to uncovering the self-immunity mechanism mediated by ScrI. While we have shed light on how ScrI protects against self-lysis, future research is needed to fully understand the exact mechanism by which ScrM recognizes target cells and cleaves the PG, as well as its potential role in fratricide among streptococci. Moreover, the efficient lytic activity of ScrM against clinically important streptococcal species such as *S. dysgalactiae, S. agalactiae* and *S. pyogenes*, highlights the potential to explore this protein for developing antimicrobial strategies targeting these species.

## Materials and methods

### Bacterial strains and growth conditions

The bacterial strains used in this work are listed in Table S5. *E. coli* strains were grown in lysogeny broth (LB) with shaking, or on lysogeny agar (LA) at 37 °C, if not stated otherwise. *S. dysgalactiae* strains were grown in airtight tubes at 37 °C in C-medium or on BHI agar at 37 °C with 10% CO_2_. Other *Streptococcus* species were grown in airtight tubes at 37 °C in C-medium or BHI. When appropriate, antibiotics were added for selection: 100 µg/ml ampicillin or 60 µg/ml spectinomycin for *E. coli*, 100 µg/ml spectinomycin for *S. dysgalactiae*. Anhydrotetracycline (aTc) was added for induction of ectopic gene expression when appropriate.

For transformation of *E. coli*, chemically competent DH5α, BL21 or IM08B cells were prepared using calcium-chloride treatment followed by transformation with heat shock according to standard protocols. *S. dysgalactiae* strains were either transformed by electroporation with plasmids isolated from *E. coli* DH5α or IM08B, or by natural transformation using linear DNA, as described previously (Mårli et al., 2024).

### DNA techniques

All plasmids used in this work are listed in Table S6. Primers used for cloning are listed in Table S7. Plasmid constructs were verified by PCR and sequencing, and knockout mutants were verified by PCR.

#### Construction of pFD116-P_scrM_-luc-gfp

To construct the pFD116-P*_scrM_-luc-gfp* reporter plasmid, the promoter region of *scrM* was amplified from genomic DNA of *S. dysgalactiae* iSDSE-NORM37 using primers mm79/mm61. The *luc-gfp* fragment was amplified using primers mm55/mm57 and genomic DNA from strain MK175 as template. The P*_cbpD_* fragment contained a 3’ end complementary to the 5’ end of the *luc-gfp* fragment, which facilitated fusion of the two fragments by overlap extension PCR (59) using primers mm79/mm57. Restriction sites at the 5’ and 3’ ends were introduced with the primers. The resulting P*_scrM_*-*luc-gfp* amplicon was cleaved with restriction enzymes SalI-HF and NheI-HF and ligated into the corresponding sites of pFD116 (60).

#### Construction of pRSET-scrM

To construct pRSET-*scrM*, the *scrM* gene encoding mature ScrM (ScrM without the signal sequence) was amplified using genomic DNA from *S. dysgalactiae* MA201 and primers ats1/ats2. DNA encoding a polyhistidine tag (6x His-tag) and a TEV-protease cleavage site was introduced with the primers at the 5’ end of *scrM*. The *scrM* amplicon was cleaved with NdeI and HindIII and ligated into pRSET-A (Invitrogen), generating pRSET-*scrM*.

#### Construction of pRSET-scrM^G302A,G303A^

To construct pRSET-*scrM^G302A,G303A^, scrM* was amplified in two fragments from pRSET-*scrM* by primer pairs pRSET(F)/jaa4 and jaa3/pRSET(R), introducing Gly to Ala mutations in position 302 and 303. The two fragments were fused by overlap extension PCR (59) using primer pair pRSET(F)/pRSET(R). The *scrM*^G302A,G303^ amplicon was cleaved with NdeI and HindIII and ligated into pRSET-A, generating pRSET-*scrM*^G302A,G303A^.

#### Construction of pRSET-sfGFP-SH3b-CCD

To construct pRSET-sfGFP-SH3b-CCD the CHAP-encoding part (amino acids 40 to 170 in the full-length protein) of *scrM* was substituted with the *sf-gfp* gene. The *sf-gfp* gene was amplified using primer pair ats3/ats4 and genomic DNA of SPH470 as template. The SH3b-CCD domains of *scrM* was amplified from *S. dysgalactiae* MA201 using primer pair ats5/ats2. The two amplicons were fused using overlap extension PCR and primer pair ats3/ats2, and subsequently cleaved with NdeI and HindIII and ligated into pRSET-A, generating pRSET-sfGFP-SH3b-CCD.

#### Construction of pRSET-sfGFP-CCD

To construct pRSET-sfGFP-CCD the *sf-gfp* gene was amplified using primer pair pRSET(F)/jaa2 and the CCD domain was amplified using primer pair jaa1/pRSET(R), both using the plasmid pRSET-sfGFP-SH3b-CCD as template. The resulting amplicons were fused together by overlap extension PCR using primer pair pRSET(F)/pRSET(R), and subsequently cleaved with NdeI and HindIII and ligated into pRSET-A, generating pRSET-sfGFP-CCD.

#### Construction of pRSET-sfGFP-CCD^,G302A,G303A^

To construct pRSET-sfGFP-CCD^G302A,G303A^ the *sf-gfp* gene was amplified using primer pair pRSET(F)/jaa2 and pRSET-sfGFP-SH3b-CCD as template. The CCD domain with G302A and G303A mutations was amplified from pRSET-*scrM*^G302A,G303A^ using primer pair jaa1/pRSET(R). The amplicons were fused together by overlap extension PCR using primer pair pRSET(F)/pRSET(R), and subsequently cleaved with NdeI and HindIII and ligated into pRSET-A, generating pRSET-sfGFP-CCD^G302A,G303A^.

#### Deletion of scrI (ΔscrI::kan)

Deletion of *scrI* was achieved using natural transformation, following the same approach as described by Mårli et al (41). The DNA cassette was constructed by overlap extension PCR fusing the ∼2000 bp region upstream and downstream of *scrI* to the 5’ and 3’ end, respectively, of a kanamycin resistance cassette. This amplicon was then transformed into *S. dysgalactiae* Stdys021 to knock out *femX2* (MM463).

#### Ectopic expression of scrI (ΔscrI::kan, pFD116-P_tet_-scrI)

To construct the expression plasmid pFD116-P_tet_-*scrI* the *scrI* gene was amplified using primer pair mm122/mm123 and *S. dysgalactiae* Stdys021 as template. The vector was amplified using primers mm120/mm121 and pFD116 as template, and the *scrI* gene was inserted into the vector behind a P_tet_ promoter using NEBuilder^®^ HiFi DNA assembly to create pFD116-P_tet_-*scrI*. The assembly reaction was transformed into *E. coli* IM08B and subsequently transformed into MM463 to give Δ*scrI::kan, pFD116-Ptet-scrI* (MM481).

### Protein expression and purification

*E. coli* BL21 containing variants of the pRSET-His-ScrM expression plasmid was grown to an OD_600_ of 0.4 at 37 °C. Protein expression was induced by adding 0.1 mM isopropyl β-D-1-thiogalactopyranoside (IPTG) followed by incubation at 25 °C for 4 h. The cells were harvested at 7,000 × *g* for 5 minutes and resuspended in 1/100 culture volume of 20 mM Tris-buffered saline (TBS) at pH 7.4 containing 500 mM NaCl and 20 mM imidazole. The cells were lysed in a French Pressure cell (SPCH-10-60, Homogenizing Systems Ltd) at 10,000 psi, and insoluble material was removed by centrifugation at 13,000 × *g* for 5 minutes followed by filtration of the supernatant using a 0.45 µm filter. His-ScrM variants were purified by Immobilized-Metal Affinity Chromatography (IMAC) using 1 ml HisTrap HP columns connected to an Äkta pure 25L (Cytiva). After binding of His-tagged proteins on the column it was washed with 10 column volumes of 20 mM Tris-HCl (pH 7.4), 500 mM NaCl, 20 mM imidazole. His-tagged proteins were eluted using 20 mM Tris-HCl (pH 7.4), 500 mM NaCl, 500 mM imidazole over a 30-ml linear gradient from 0 – 100% elution buffer. Proteins were detected at 280 nm. To remove imidazole and excess salt, eluted protein was dialyzed against 10 mM Tris-HCl with 150 mM NaCl for 1 h at room temperature. Proteins were analyzed by SDS-PAGE, and protein concentration was estimated by Abs280 measurements (Nanodrop1000, Thermo Fisher Scientific).

### Luciferase reporter assay

Overnight cultures of reporter strains were initially diluted at a 1/50 ratio in C-medium in airtight tubes and incubated at 37 °C until mid-logarithmic phase. The cultures were subsequently diluted to an OD_600_ of ∼0.03 in C-medium containing a final concentration of 170 µg/ml D-luciferin (Invitrogen™). Volumes of 300 µl culture were added in triplicate in white clear-bottom 96-well plates (Corning), and OD_600_ and luminescence were measured at 5- or 10-minute intervals in a Hidex Sense microplate reader at 37 °C. When appropriate, XIP2 was added to a final concentration of 250 ng/ml when OD_600_ reached ∼0.10-0.15. As a control for background levels, a 300 µl sample of the cell-free growth medium containing D-luciferin was used.

### Zymogram analysis of murein hydrolase activity

Zymogram analysis was carried out as described by Eldholm et al. (26) with some modifications. In brief, 15 µl of whole cell extracts from both competent and non-competent cells were loaded onto the SDS-PAGE gel, alongside purified protein (1 µg, 2.5 µg and 5 µg). The samples were separated by SDS-PAGE using a 4% stacking gel and a 12% resolving gel containing heat-treated cells (95 °C, 10 minutes) of *S. dysgalactiae* Stdys021. Following electrophoresis at 2.5 V cm^-2^, the gels were washed in de-ionized water for 2 x 30 minutes before adding the refolding buffer (50 mM NaCl, 20 mM MgCl2, 0.5% Triton X-100, and 20 mM Tris-HCl, pH 7.4). The gel was incubated in refolding buffer until lytic zones could be observed.

For the preparation of whole cell extracts, cells were grown to an OD_600_ of 0.1. To induce competence, 250 ng/ml XIP2 was added to one culture, and both cultures were incubated for 1 hour before harvesting by centrifugation at 4000 x *g* for 10 minutes.

### Isolation of peptidoglycan

Strains MM358 and MM463 were inoculated in 1 L C-medium, while strain MM481 was inoculated in 1 L C-medium containing 100 ng/ml aTc for expression of *scrI* and 100 µg/ml spectinomycin for selection. At an OD_600_ of 0.2, 250 ng/ml XIP2 was added to induce cells to competence. Cultures were incubated further for 1 hour and cells were harvested at 10,000 × *g* for 10 minutes. Peptidoglycan (PG) was purified as previously described by Vollmer (61). The isolated PG was lyophilized and resuspended in water to a final concentration of 25 mg/ml.

### LC-MS/MS

Two mg of purified PG were treated with 1 ml 48% hydrofluoric acid (HF) at room temperature for 16 hours with gentle mixing. The HF treated PG was collected by centrifugation at 20,000 × *g* for 5 minutes and washed four times with 1 ml dH_2_O. The TA-free PG was lyophilized and dissolved in dH_2_O to a final concentration of 25 mg/ml.

Next, 1 mg PG was digested overnight at 37 °C with 40 µg/ml mutanolysin in 100 µl 50 mM sodium phosphate buffer (pH 6). After mutanolysin treatment, the enzyme was inactivated and precipitated by incubating the samples at 95 °C for 10 minutes, and insoluble material was removed by centrifugation at 20,000 × *g* for 10 minutes. The supernatant was added to an equal volume of 0.5 M boric acid (pH 9) and treated with 1-2 mg sodium borohydride for 30 minutes at room temperature to reduce the sugars. The reaction was stopped by adjusting the pH to 2.0-3.0 using 20% phosphoric acid.

For purification of the digested PG, the sample was applied to SPE-tips with 1.5 mg Supelclean™ENVI-Carb™. The tips were first conditioned with 2 x 150 µl 100% acetonitrile (ACN), followed by equilibration with 2 x 150 µl of 0.5% trifluoroacetic acid (TFA). Following this, 50 µl of the acidified sample was loaded onto the tips and washed twice with 150 µl 0.5% TFA. The samples were then eluted with 2 x 40 µl of 70% CAN. Centrifugation was performed at 1500 x *g* for 3 minutes. The eluted samples were dired using a Speed-Vas and resuspended in 0.1% formic acid for LC-MS/MS analysis.

The samples were analyzed on a an Ultimate RSLCnano/QExactive (Thermo Fisher Scientific) system. The dried samples were dissolved in solution A (.1 % formic acid, 2% ACN) and injected directly onto a 50 cmx75 µm analytical column (Acclaim PepMap RSLC C18, 2 µm, 100 Å, 75 µm i.d. x 50 cm, nanoViper). A 60 min method was used for chromatographic separation at a flow rate of 300 nl/min.: from 2 to 10 % solution B (99.9 % ACN, 0.1% formic acid) over 35 min, followed by an increase to 25 % B over 5 min, a wash step of 80% B over 5 min, and finally an equilibration step at 2% B of 15 min.

The Q-Exactive mass spectrometer was set up as follows (Top3 method): a full scan (150-2250 m/z) at R=70.000 was followed by 3 MS2 scans at R=17500, using an NCE setting of 28. AGC was set at 1E6 at MS1 level, and 1E5 at MS2 level. Precursors with z>5 were excluded, but singly charged precursors were allowed. Dynamic exclusion was set to 20 seconds.

### Muropeptide analysis

For automated analysis of muropeptides, we generated an *in silico* MS/MS library using PGN_MS2 (62), including variations in glycans (GlcNAc, MurNAc, and O-acetylated MurNAc), stem peptides (L-Ala-D-(γ)-iGln-L-Lys-D-Ala-D-Ala), and interpeptide bridges (Gly, Ala, Ser, Thr). MS-DIAL (v.5) was used to process the raw LC-MS/MS data, compare it to the *in-silico* library, and identify muropeptides based on *m/z*, MS1 isotopic patterns, and MS/MS spectral similarity. Muropeptides with MS-DIAL scores >1 was considered identified.

For comparative analysis, muropeptides identified by MS-DIAL in the Δ*scrI* mutant and the ScrI expression strain were visualized and quantified using the software Freestyle (v1.4; Thermo Fisher Scientific). Selected muropeptide-pairs of 30 Da mass difference was compared via extracted-ion chromatograms, with intensities normalized to the most-intense peak. Manual *de novo* annotation of MS/MS-spectra was performed by comparing mass differences of the most prominent peaks to identify fragment ion series indicative of the muropeptide stem- and bridge amino acid sequence. The manual findings (b-/y-fragmentation series or ion-series displaying 30 Da differences) were validated by comparing the *m/z*-values to theoretical MS/MS-spectra predicted by PGN_MS2 and identified with MS-DIAL to obtain peptide structures.

### Phase contrast and fluorescence microscopy

For microscopy analyses of whole cells, overnight cultures were diluted to an OD_600_ of 0.05 in C-medium and grown to an OD_600_ of ∼0.2. When appropriate, 250 ng/ml XIP2 was added to induce competence at an OD_600_ of ∼0.1 and the cells were incubated further for 20 minutes before harvesting the cells at 4,000 x *g* for 10 minutes. The cells were then washed once with 1X PBS and stained with 15 µg/ml of various sfGFP-ScrM fusion protein constructs in PBS-Tween in the dark, at room temperature for 8 minutes. After staining, the cells were washed three times in 1X PBS and immobilized on agarose pads (1.2%) before imaging on a Zeiss AxioObserver with ZEN Blue software. Images were captured with an ORCA-Flash4.0 V2 Digital CMOS camera (Hamamatsu Photonics) through a 100x PC objective. For fluorescence microscopy, HPX 120 Illuminator (Zeiss) was used as a light source. Image analysis was performed using MicrobeJ (63).

### ScrM sensitivity assays

For examining sensitivity to ScrM in liquid cultures, overnight cultures of *streptococcus* species were diluted 1/50 in C-medium or BHI and grown to an OD_600_ of 0.4-0.6. Cultures were then diluted to an OD_600_ of ∼0.03 and growth of cultures (300 µl in a flat-bottom 96-well plate (Sarstedt)) were monitored at 37 °C in a Hidex Sense microplate reader at 10-minute intervals. When appropriate, cells were induced to competence by addition of 250 ng/ml XIP2 at an OD_600_ of ∼0.10. Purified ScrM (1 µg/ml or 10 µg/ml) was added 20 minutes post XIP2 induction, or at an OD_600_ of ∼ 0.15-0.20. When appropriate ScrM inactivated by treatment with DTT (150 mM) for 5 minutes followed by iodoacetamide (0.5 mM) treatment for 15 minutes was used as a negative control (64).

## Data availability

All data in the study are included in the article or in Supporting information.

## Acknowledgements

We thank the Norwegian Veterinary Institute for kindly providing the *Streptococcus suis* strain used in this work. We thank Davide Porcellato (Norwegian University of Life Sciences) for help with *S. dysgalactiae* strains. We also thank Morten Skaugen (Norwegian University of Life Sciences) and Per Kristian Edvardsen (Norwegian University of Life Sciences) for technical assistance with LC-MS. Mass spectrometry-based proteomic analyses were performed by the MS and Proteomics Core Facility, Norwegian University of Life Sciences (NMBU). This facility is a member of the National Network of Advanced Proteomics Infrastructure (NAPI), which is funded by the Research Council of Norway INFRASTRUKTUR-program (project number: 295910). MK was supported by a JPI-AMR grant from the Research Council of Norway (Grant No. 296906). MA was supported by the Novo Nordisk Foundation through grant NNF20OC0061313.

## Supporting information titles and legends

**Fig. S1.** SDS-PAGE analysis of ScrM and its fusion protein variants expressed in *E. coli* and purified using Immobilized-Metal Affinity Chromatography (IMAC). The proteins were separated on a 12% SDS-PAGE gel and stained with Coomassie Brilliant Blue. Lanes 1-6: purified proteins corresponding to the constructs shown in the schematic on the right, representing different ScrM constructs with respective domains: CHAP domain, SH3b domain and a C-terminal conserved domain (CCD). The CCD^mut^ variants harbors G302A and G303A substitutions.

**Fig. S2.** Fluorescence microscopy images showing the binding of sfGFP-SH3b-CCD on *S. pyogenes* (top row) and *S. pneumoniae* (bottom row). Scale bars represent 2 µm.

**Fig. S3. (A)** Multiple sequence alignment of the conserved C-terminal domain (CCD) across streptococcal species, including *S. dysgalactiae* (MA201-1_01842), *S. thermophilus* (AKH33394.1), *S. vestibularis* (WP_117648089), *S. ictaluri* (WP_008089638.1)*, S. pyogenes* (AAK33168), *S. equi* (WP_012677198.1)*, S. porcinus* (WP_003083717), *S. uberis* (WP_203261572.1) and *S. iniae* (WP_071127360). The alignment highlights conserved residues within the CCD domain, with fully conserved residues marked with an asterisk (*), residues with strongly similar properties indicated by a colon (:), and residues with weakly similar properties marked by a period (.). The LAGG motif, which is conserved across all species, is emphasized with a black box. **(B)** AlphaFold 3 structural prediction of ScrM. The structure prediction is color-coded based on model confidence, where dark blue represents the highest confidence (pLDDT > 90) and red represents the lowest confidence (pLDDT < 50). **(C)** AlphaFold Structural prediction of the conserved C-terminal domain (CCD). The conserved LAGG motif is highlighted in yellow sticks, with the amino acids and positions noted. **(D)** Microscopy of sfGFP-CCD^mut^ localization and heatmap analysis. Localization of the mutant sfGFP-CCD^mut^ protein where the LAGG motif is mutated to LAAA (G302A,G303A) on *S. dysgalactiae* cells. The heatmap analysis (n = 1060) indicates the distribution of sfGFP-CCD^mut^ across the cell population. **(E)** Growth curves of *S. dysgalactiae* with addition of ScrM^WT^, ScrM with mutated LAGG to LAAA motif (ScrM^mut^) and ScrM inactivated by treatment with 50 mM iodoacetamide (ScrM^IAM^). Growth was monitored by measuring OD_600_ at 10-min intervals. ScrM was added to cultures at an OD_600_ of ∼ 0.2. The data represent the mean of three technical replicates, with error bars indicating standard deviation.

**Fig. S4.** Complementation of ScrI. Growth curves of *S. dysgalactiae* Δ*scrI* with ectopic expression of *scrI* from a tetracycline inducible promoter (MM481) in the presence of 0 ng/ml, 50 ng/ml or 100 ng/ml aTc. A final concentration of 1 µg/ml ScrM was added at OD_600_ = 0.2. OD_600_ was measured at 10-minute intervals An untreated control was included. Data represent the mean of three technical replicates, with error bars indicating standard deviation. The results shown are representative of three independent experiments.

**Fig. S5. (A**) LC-MS/MS chromatograms showing the comparison of muropeptide peaks in competent cells of the Δ*scrI* mutant (top, strain MM463) and the Δ*scrI* mutant ectopically expressing ScrI (bottom, strain MM481). The Δ*scrI* mutant displays a peak at *m/z* = 555 (*z* = 2), while the strain expressing ScrI shows a 10-fold reduction of this peak but instead an additional peak at *m/z* = 570 (*z* =2), indicating a total 30 Da increase in mass. Notably, this peak is not present in the Δ*scrI* mutant **(B)** LC-MS/MS spectra for *de novo* analysis of mass peaks *m/z* = 555 in the competent Δ*scrI* mutant (top, strain MM463) highlighting signature ions representing a fragment of the stem peptide without a bridge (*m/z* = 676) and the same fragment with increasing bridge components (*m/z* = 747, 818, 889), verifying the AAA bridge. A second signature ion is the *m/z* = 431, which represent a stem-fragment with AK plus the entire bridge. In the competent ScrI expression strain (bottom, strain MM481) we see the same stem fragment without bridge (*m/z* = 676), but a shifted series of +30 Da compared to the Δ*scrI* mutant, and it is clear that the + 30 Da shift occurs in position one indicating an Ala➔Thr modification. Similarly, the stem-fragment with AK plus the entire bridge is now 30 Da heavier. *m/z* = 555 was determined to be AqKA-[3-NH2-AAA], while *m/z* = 570 was determined to be (AqKA-[3-NH2-TAA]. A: Alanine, q: γ-iso-Glutamine; K: Lysine.

**Table S1** Characteristics of *Streptococcus* species tested for sensitivity to ScrM, including interpeptide bridges, cell wall polysaccharides and fratricins and their domains

**Table S2.** Top 4 predicted reduced muropeptides by PGN_MS2 and MS-DIAL in *S. dysgalactiae* Δ*scrI*.

**Table S3.** Top 4 predicted reduced muropeptides by PGN_MS2 and MS-DIAL in *S. dysgalactiae* Δ*scrip*, Ptet-*scrI*.

**Table S4.** Top 4 predicted reduced muropeptides by PGN_MS2 and MS-DIAL in *S. dysgalactiae* WT.

**Table S5.** Strains used in this study.

**Table S6.** Plasmids used in this study.

**Table S7.** Oligos used in this study.

**Text S1.** DNA sequences encoding *scrM* and *scrI*.

## References

1. W. Vollmer, B. Joris, P. Charlier, S. Foster, Bacterial peptidoglycan (murein) hydrolases. FEMS Microbiol Rev 32, 259–286 (2008).

2. R. Young, I.-N. Wang, W. D. Roof, Phages will out: strategies of host cell lysis. Trends Microbiol 8, 120–128 (2000).

3. O. Johnsborg, V. Eldholm, M. L. Bjørnstad, L. S. Håvarstein, A predatory mechanism dramatically increases the efficiency of lateral gene transfer in *Streptococcus pneumoniae* and related commensal species. Mol Microbiol 69, 245–253 (2008).

4. R. Dziarski, D. Gupta, The peptidoglycan recognition proteins (PGRPs). Genome Biol 7, 232 (2006).

5. W. Vollmer, D. Blanot, M. A. De Pedro, Peptidoglycan structure and architecture. FEMS Microbiol Rev 32, 149–167 (2008).

6. E. Sauvage, F. Kerff, M. Terrak, J. A. Ayala, P. Charlier, The penicillin-binding proteins: structure and role in peptidoglycan biosynthesis. FEMS Microbiol Rev 32, 234–258 (2008).

7. K. H. Schleifer, O. Kandler, Peptidoglycan types of bacterial cell walls and their taxonomic implications. Bacteriol Rev 36, 407–477 (1972).

8. S. Rohrer, B. Berger-Bächi, FemABX peptidyl transferases: a link between branched-chain cell wall peptide formation and beta-lactam resistance in gram-positive cocci. Antimicrob Agents Chemother 47, 837–846 (2003).

9. A. Fiser, S. R. Filipe, A. Tomasz, Cell wall branches, penicillin resistance and the secrets of the MurM protein. Trends Microbiol 11, 547–553 (2003).

10. A. Vermassen et al., Cell wall hydrolases in bacteria: insight on the diversity of cell wall amidases, glycosidases and peptidases toward peptidoglycan. Front Microbiol 10, 331 (2019).

11. L. Fontaine, A. Wahl, M. Fléchard, J. Mignolet, P. Hols, Regulation of competence for natural transformation in streptococci. Infect Genet Evol 33, 343–360 (2015).

12. L. Mashburn-Warren, D. A. Morrison, M. J. Federle, A novel double-tryptophan peptide pheromone controls competence in *Streptococcus* spp. via an Rgg regulator. Mol Microbiol 78, 589–606 (2010).

13. D. A. Morrison, E. Guédon, P. Renault, Competence for natural genetic transformation in the *Streptococcus bovis* group streptococci *S. infantarius* and *S. macedonicus*. J Bacteriol 195, 2612–2620 (2013).

14. L. Fontaine et al., A novel pheromone quorum-sensing system controls the development of natural competence in *Streptococcus thermophilus* and *Streptococcus salivarius*. J Bacteriol 192, 1444–1454 (2010).

15. E. Zaccaria et al., Control of competence for DNA transformation in *Streptococcus suis* by genetically transferable pherotypes. PLoS One 9, e99394 (2014).

16. C. Johnston, B. Martin, G. Fichant, P. Polard, J. P. Claverys, Bacterial transformation: distribution, shared mechanisms and divergent control. Nat Rev Microbiol 12, 181–196 (2014).

17. O. Xie et al., Inter-species gene flow drives ongoing evolution of *Streptococcus pyogenes* and *Streptococcus* dysgalactiae subsp. *equisimilis*. Nat Commun 15, 2286 (2024).

18. N. J. Croucher et al., Rapid pneumococcal evolution in response to clinical interventions. Science 331, 430–434 (2011).

19. C. Chewapreecha et al., Dense genomic sampling identifies highways of pneumococcal recombination. Nature Genetics 46, 305–309 (2014).

20. D. Straume, G. A. Stamsås, L. S. Håvarstein, Natural transformation and genome evolution in *Streptococcus pneumoniae*. Infect Genet Evol 33, 371–380 (2015).

21. C. Johnston, N. Campo, M. J. Bergé, P. Polard, J.-P. Claverys, *Streptococcus pneumoniae*, le transformiste. Trends Microbiol 22, 113–119 (2014).

22. G. Salvadori, R. Junges, D. A. Morrison, F. C. Petersen, Competence in *Streptococcus pneumoniae* and close commensal relatives: mechanisms and implications. Front Cell Infect Microbiol 9, 94 (2019).

23. J. P. Claverys, B. Martin, L. S. Håvarstein, Competence-induced fratricide in streptococci. Mol Microbiol 64, 1423–1433 (2007).

24. L. Kausmally, O. Johnsborg, M. Lunde, E. Knutsen, L. S. Håvarstein, Choline-binding protein D (CbpD) in *Streptococcus pneumoniae* is essential for competence-induced cell lysis. J Bacteriol 187, 4338–4345 (2005).

25. K. H. Berg, T. J. Biørnstad, O. Johnsborg, L. S. Håvarstein, Properties and biological role of streptococcal fratricins. Appl Environ Microbiol 78, 3515–3522 (2012).

26. V. Eldholm et al., Pneumococcal CbpD is a murein hydrolase that requires a dual cell envelope binding specificity to kill target cells during fratricide. Mol Microbiol 76, 905–917 (2010).

27. Y. Zhu et al., CrfP, a fratricide protein, contributes to natural transformation in *Streptococcus suis*. Vet Res 52, 50 (2021).

28. N. Cullin, S. Redanz, K. J. Lampi, J. Merritt, J. Kreth, Murein Hydrolase LytF of *Streptococcus sanguinis* and the Ecological Consequences of Competence Development. Appl Environ Microbiol 83 (2017).

29. K. H. Berg, H. S. Ohnstad, L. S. Håvarstein, LytF, a novel competence-regulated murein hydrolase in the genus *Streptococcus*. J Bacteriol 194, 627–635 (2012).

30. S. R. Gargis et al., Use of 4-sulfophenyl isothiocyanate labeling and mass spectrometry to determine the site of action of the streptococcolytic peptidoglycan hydrolase zoocin A. Appl Environ Microbiol 75, 72–77 (2009).

31. L. S. Heath et al., The streptococcolytic enzyme zoocin A is a penicillin-binding protein. FEMS Microbiol Letters 236, 205–211 (2004).

32. J. R. Vidal Amaral et al., Bacteriocin Producing *Streptococcus agalactiae* Strains Isolated from Bovine Mastitis in Brazil. Microorganisms 10, 588 (2022).

33. J. Pei, D. A. Mitchell, J. E. Dixon, N. V. Grishin, Expansion of type II CAAX proteases reveals evolutionary origin of γ-secretase subunit APH-1. J Mol Biol 410, 18–26 (2011).

34. L. S. Håvarstein, B. Martin, O. Johnsborg, C. Granadel, J.-P. Claverys, New insights into the pneumococcal fratricide: relationship to clumping and identification of a novel immunity factor. Mol Microbiol 59, 1297–1037 (2006).

35. D. Straume, G. A. Stamsås, Z. Salehian, L. S. Håvarstein, Overexpression of the fratricide immunity protein ComM leads to growth inhibition and morphological abnormalities in *Streptococcus pneumoniae*. Microbiology 163, 9–21 (2017).

36. M. J. Bergé et al., A programmed cell division delay preserves genome integrity during natural genetic transformation in *Streptococcus pneumoniae*. Nat Commun 8, 1621 (2017).

37. S. R. Gargis et al., Zif, the Zoocin A Immunity Factor, Is a FemABX-Like Immunity Protein with a Novel Mode of Action. Appl Environ Microbiol 75, 6205–6210 (2009).

38. S. A. Beatson, G. L. Sloan, R. S. Simmonds, Zoocin A immunity factor: a femA-like gene found in a group C *Streptococcus*. FEMS Microbiol Lett 163, 73–77 (1998).

39. S. Brouwer et al., Pathogenesis, epidemiology and control of Group A *Streptococcus* infection. Nat RevMicrobiol 21, 431–447 (2023).

40. A. Jensen, M. Kilian, Delineation of *Streptococcus dysgalactiae*, its subspecies, and its clinical and phylogenetic relationship to *Streptococcus pyogenes*. J Clin Microbiol 50, 113–126 (2012).

41. M. T. Mårli, O. Oppegaard, D. Porcellato, D. Straume, M. Kjos, Genetic modification of *Streptococcus dysgalactiae* by natural transformation. mSphere 0, e00214–00224 (2024).

42. P. Mitkowski et al., Structural bases of peptidoglycan recognition by lysostaphin SH3b domain. Sci Rep 9, 5965 (2019).

43. E. Tamai et al., X-ray structure of a novel endolysin encoded by episomal phage phiSM101 of *Clostridium perfringens*. Mol Microbiol 92, 326–337 (2014).

44. J. Z. Lu, T. Fujiwara, H. Komatsuzawa, M. Sugai, J. Sakon, Cell wall-targeting domain of glycylglycine endopeptidase distinguishes among peptidoglycan cross-bridges. J Biol Chem 281, 549–558 (2006).

45. C. J. Porter et al., The 1.6 A crystal structure of the catalytic domain of PlyB, a bacteriophage lysin active against *Bacillus anthracis*. J Mol Biol 366, 540–550 (2007).

46. H. Guérin, S. Kulakauskas, M. P. Chapot-Chartier, Structural variations and roles of rhamnose-rich cell wall polysaccharides in Gram-positive bacteria. J Biol Chem 298, 102488 (2022).

47. M. Schlag et al., Role of staphylococcal wall teichoic acid in targeting the major autolysin Atl. Mol Microbiol 75, 864–873 (2010).

48. S. R. Filipe, M. G. Pinho, A. Tomasz, Characterization of the *murMN* operon involved in the synthesis of branched peptidoglycan peptides in *Streptococcus pneumoniae*. J Biol Chem 275, 27768–27774 (2000).

49. S. R. Filipe, E. Severina, A. Tomasz, Functional analysis of *Streptococcus pneumoniae* MurM reveals the region responsible for its specificity in the synthesis of branched cell wall peptides. J Biol Chem 276, 39618–39628 (2001).

50. M. Beukes, J. W. Hastings, Self-protection against cell wall hydrolysis in *Streptococcus milleri* NMSCC 061 and analysis of the millericin B operon. Appl Environ Microbiol 67, 3888–3896 (2001).

51. L. S. Heath et al., Plasmid-specified FemABX-like immunity factor in *Staphylococcus sciuri* DD 4747. FEMS Microbiol Lett 249, 227–231 (2005).

52. H. P. DeHart, H. E. Heath, L. S. Heath, P. A. LeBlanc, G. L. Sloan, The lysostaphin endopeptidase resistance gene (*epr*) specifies modification of peptidoglycan cross bridges in *Staphylococcus simulans* and *Staphylococcus aureus*. Appl Environ Microbiol 61, 1475–1479 (1995).

53. M. Sugai et al., *epr*, which encodes glycylglycine endopeptidase resistance, is homologous to *femAB* and affects serine content of peptidoglycan cross bridges in *Staphylococcus capitis* and *Staphylococcus aureus*. J Bacteriol 179, 4311–4318 (1997).

54. S. Willing, E. Dyer, O. Schneewind, D. Missiakas, FmhA and FmhC of *Staphylococcus aureus* incorporate serine residues into peptidoglycan cross-bridges. J Biol Chem 295, 13664–13676 (2020).

55. A. Bateman, N. D. Rawlings, The CHAP domain: a large family of amidases including GSP amidase and peptidoglycan hydrolases. Trends Biochem Sci 28, 234–237 (2003).

56. D. Straume et al., Class A PBPs have a distinct and unique role in the construction of the pneumococcal cell wall. Proc Natl Acad Sci U S A 117, 6129–6138 (2020).

57. P. J. Moynihan, A. J. Clarke, O-Acetylated peptidoglycan: controlling the activity of bacterial autolysins and lytic enzymes of innate immune systems. Int J Biochem Cell Biol 43, 1655–1659 (2011).

58. J. Bonnet et al., Peptidoglycan O-acetylation is functionally related to cell wall biosynthesis and cell division in *Streptococcus pneumoniae*. Mol Microbiol 106, 832–846 (2017).

59. R. Higuchi, B. Krummel, R. K. Saiki, A general method of in vitro preparation and specific mutagenesis of DNA fragments: study of protein and DNA interactions. Nucleic Acids Res 16, 7351–7367 (1988).

60. F. Depardieu, D. Bikard, Gene silencing with CRISPRi in bacteria and optimization of dCas9 expression levels. Methods 172, 61–75 (2020).

61. W. Vollmer, Preparation and analysis of pneumococcal murein (peptidoglycan). Molecular Biology of Streptococci (2007).

62. J. M. C. Kwan et al., In silico MS/MS prediction for peptidoglycan profiling uncovers novel anti-inflammatory peptidoglycan fragments of the gut microbiota. Chem Sci 15, 1846–1859 (2024).

63. A. Ducret, E. M. Quardokus, Y. V. Brun, MicrobeJ, a tool for high throughput bacterial cell detection and quantitative analysis. Nat Microbiol 1, 16077 (2016).

64. M. Alcorlo, S. Martínez-Caballero, R. Molina, J. A. Hermoso, Carbohydrate recognition and lysis by bacterial peptidoglycan hydrolases. Curr Opin Struct Biol 44, 87–100 (2017).

