## Supplementary material for "Self-immunity towards a novel competence-induced streptococcal murein hydrolase is mediated by a Fem-transferase-like protein": Supplmental information

### Supporting figures

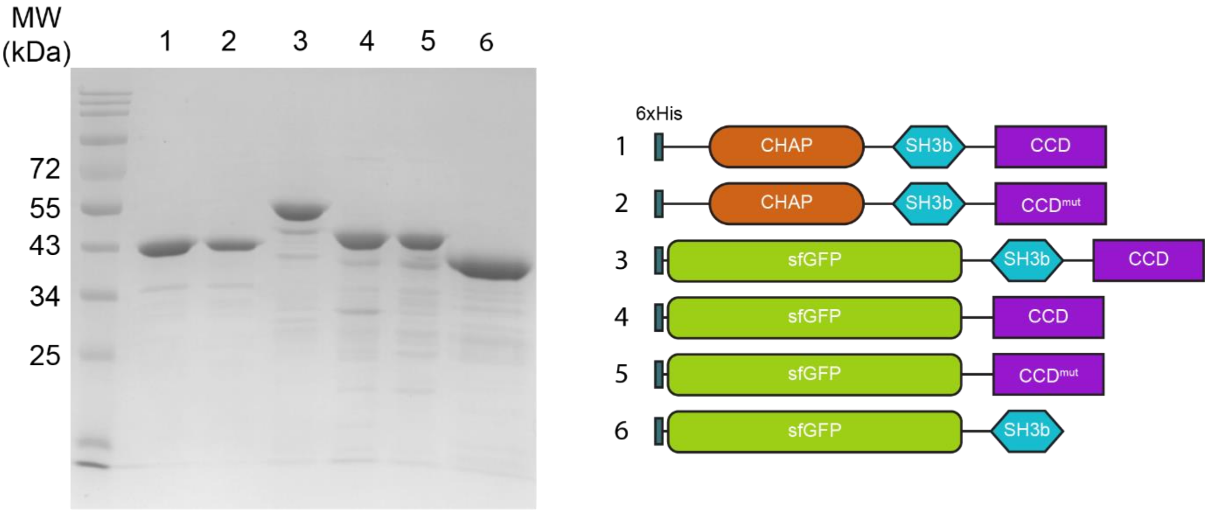


**Fig. S1.** SDS-PAGE analysis of ScrM and its fusion protein variants expressed in *E. coli* and purified using Immobilized-Metal Affinity Chromatography (IMAC). The proteins were separated on a 12% SDS-PAGE gel and stained with Coomassie Brilliant Blue. Lanes 1-6: purified proteins corresponding to the constructs shown in the schematic on the right, representing different ScrM constructs with respective domains: CHAP domain, SH3b domain and a C-terminal conserved domain (CCD). The CCD$^{mut}$ variants harbors G302A and G303A substitutions.

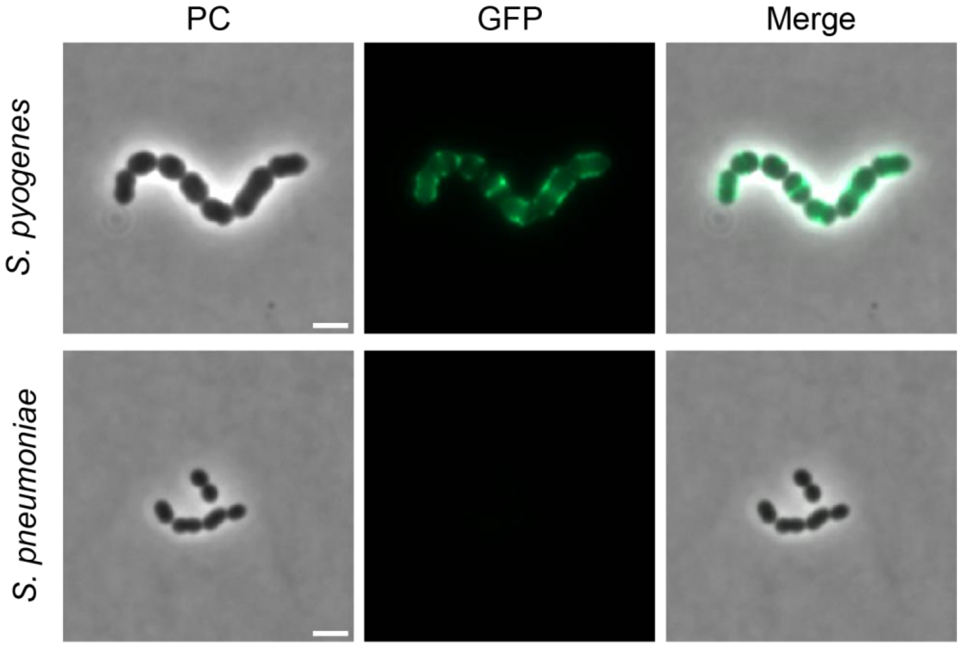


**Fig. S2.** Fluorescence microscopy images showing the binding of sfGFP-SH3b-CCD on *S. pyogenes* (top row) and *S. pneumoniae* (bottom row). Scale bars represent 2 µm.

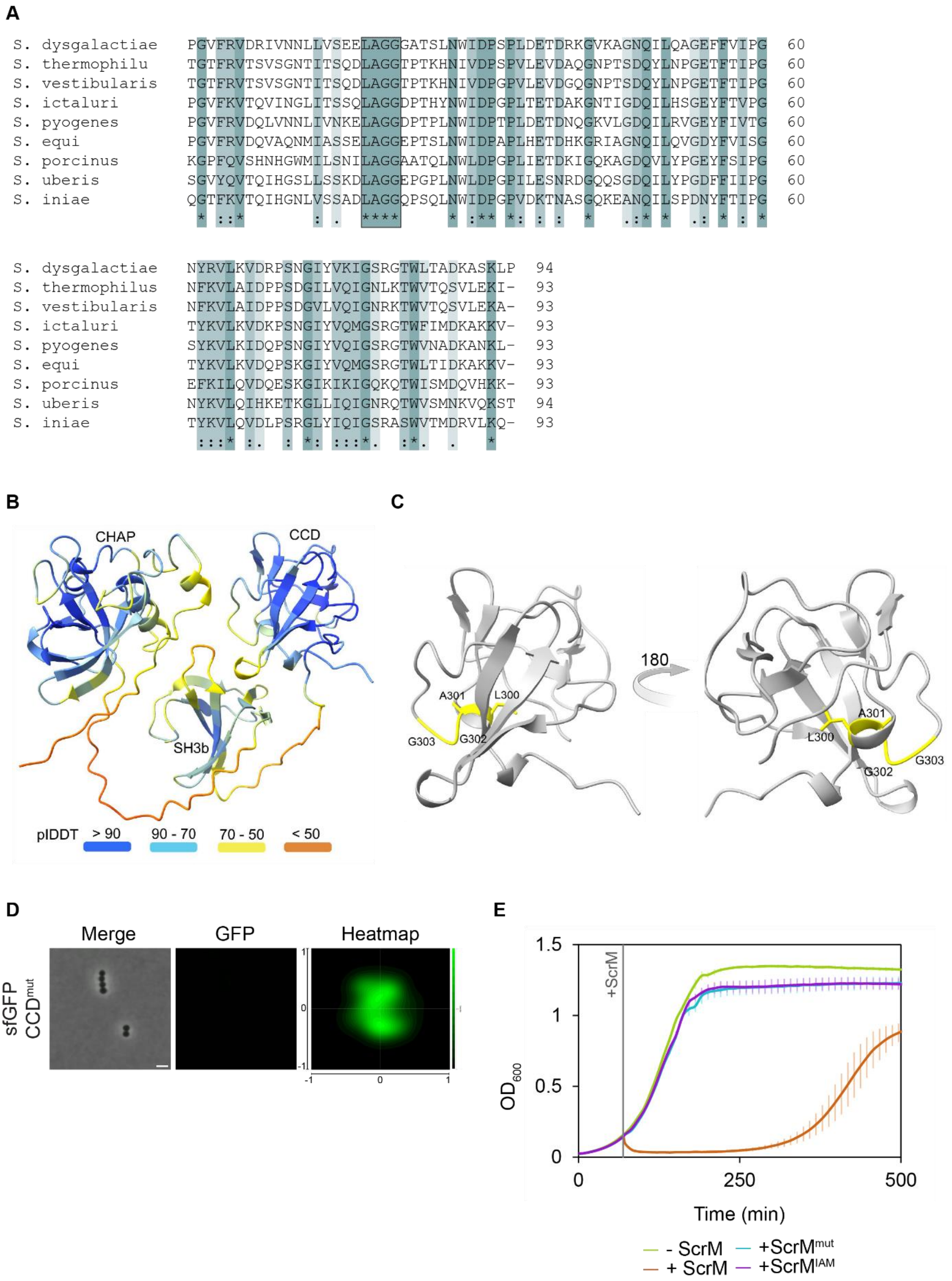


**Fig. S3. (A)** Multiple sequence alignment of the conserved C-terminal domain (CCD) across streptococcal species, including *S. dysgalactiae* (MA201-1_01842), *S. thermophilus*

(AKH33394.1), *S. vestibularis* (WP_117648089), *S. ictaluri* (WP_008089638.1), *S. pyogenes* (AAK33168), *S. equi* (WP_012677198.1), *S. porcinus* (WP_003083717), *S. uberis* (WP_203261572.1) and *S. iniae* (WP_071127360). The alignment highlights conserved residues within the CCD domain, with fully conserved residues marked with an asterisk (*), residues with strongly similar properties indicated by a colon (:), and residues with weakly similar properties marked by a period (.). The LAGG motif, which is conserved across all species, is emphasized with a black box. **(B)** AlphaFold 3 structural prediction of ScrM. The structure prediction is color-coded based on model confidence, where dark blue represents the highest confidence (pLDDT > 90) and red represents the lowest confidence (pLDDT < 50). **(C)** AlphaFold Structural prediction of the conserved C-terminal domain (CCD). The conserved LAGG motif is highlighted in yellow sticks, with the amino acids and positions noted. **(D)** Microscopy of sfGFP-CCD$^{mut}$ localization and heatmap analysis. Localization of the mutant sfGFP-CCD$^{mut}$ protein where the LAGG motif is mutated to LAAA (G302A,G303A) on *S. dysgalactiae* cells. The heatmap analysis (n = 1060) indicates the distribution of sfGFP-CCD$^{mut}$ across the cell population. **(E)** Growth curves of *S. dysgalactiae* with addition of ScrM$^{WT}$, ScrM with mutated LAGG to LAAA motif (ScrM$^{mut}$) and ScrM inactivated by treatment with 50 mM iodoacetamide (ScrM$^{IAM}$). Growth was monitored by measuring $OD_{600}$ at 10-min intervals. ScrM was added to cultures at an $OD_{600}$ of ~ 0.2. The data represent the mean of three technical replicates, with error bars indicating standard deviation.

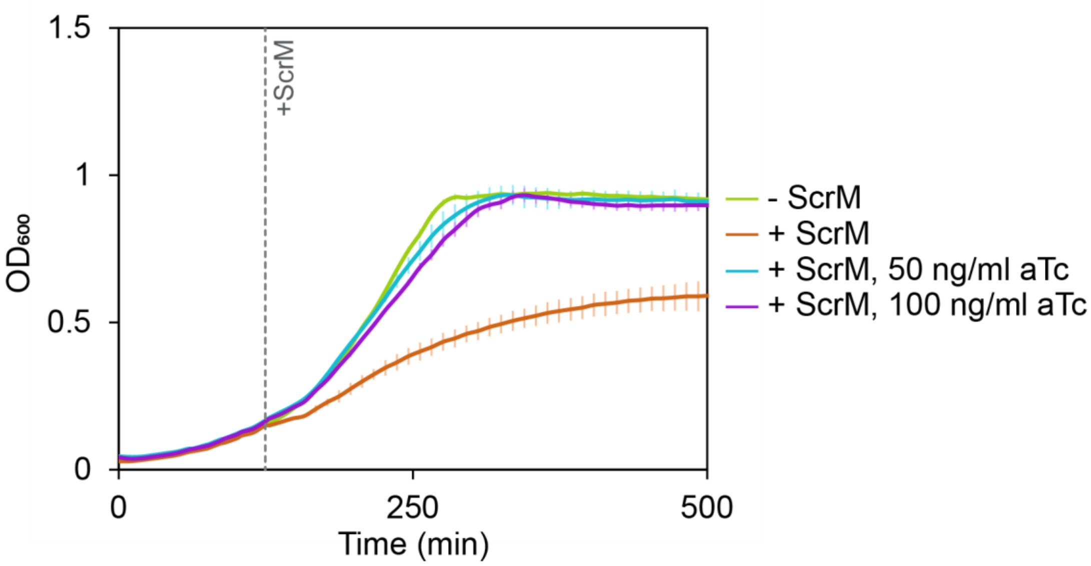


**Fig. S4**. Complementation of ScrI. Growth curves of *S. dysgalactiae* Δ*scrI* with ectopic expression of *scrI* from a tetracycline inducible promoter (MM481) in the presence of 0 ng/ml, 50 ng/ml or 100 ng/ml aTc. A final concentration of 1 µg/ml ScrM was added at $OD_{600}$ = 0.2. $OD_{600}$ was measured at 10-minute intervals An untreated control was included. Data represent the mean of three technical replicates, with error bars indicating standard deviation. The results shown are representative of three independent experiments.

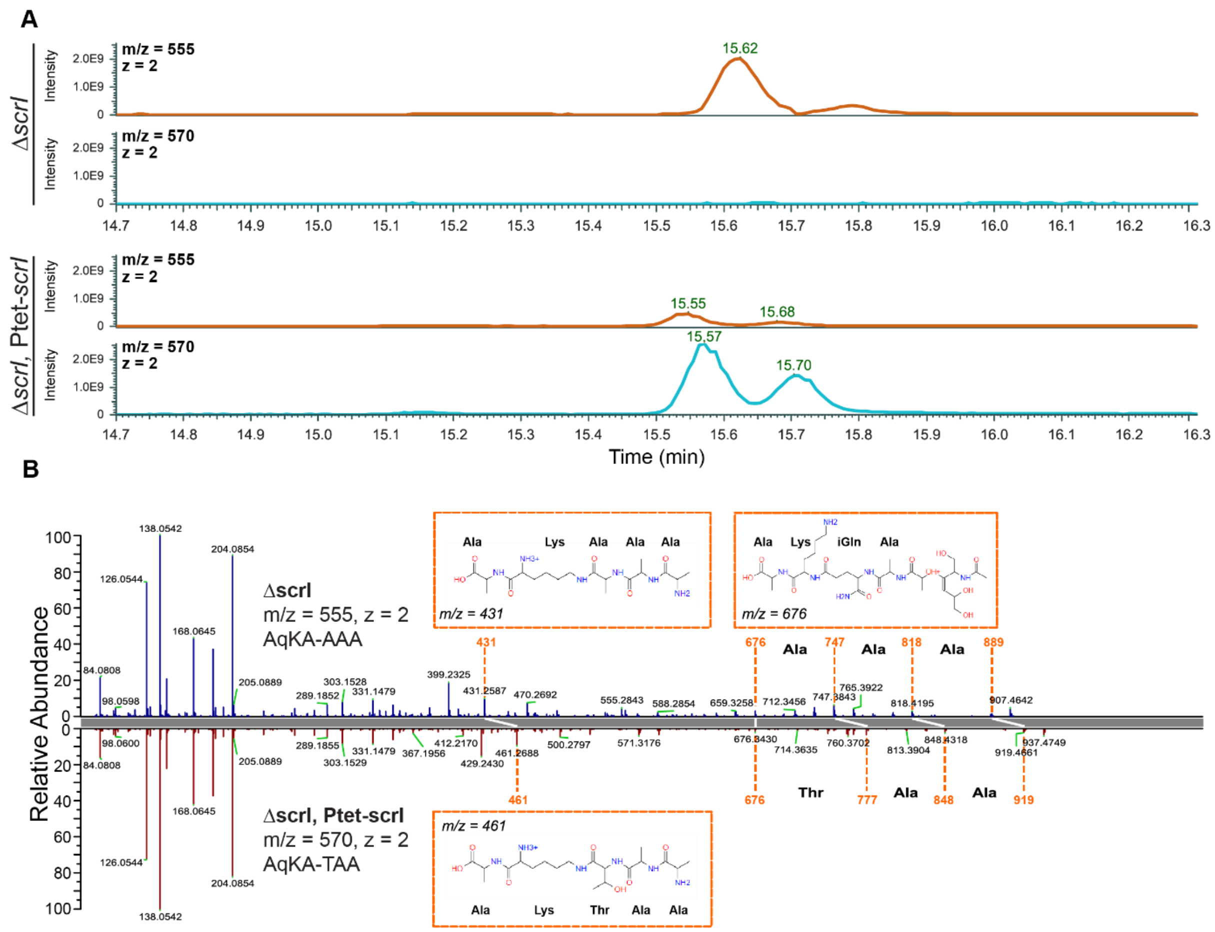


**Fig. S5. (A)** LC-MS/MS chromatograms showing the comparison of muropeptide peaks in competent cells of the Δ*scrI* mutant (top, strain MM463) and the Δ*scrI* mutant ectopically expressing ScrI (bottom, strain MM481). The Δ*scrI* mutant displays a peak at *m/z* = 555 (*z* = 2), while the strain expressing ScrI shows a 10-fold reduction of this peak but instead an additional peak at *m/z* = 570 (*z* =2), indicating a total 30 Da increase in mass. Notably, this peak is not present in the Δ*scrI* mutant **(B)** LC-MS/MS spectra for *de novo* analysis of mass peaks *m/z* = 555 in the competent Δ*scrI* mutant (top, strain MM463) highlighting signature ions representing a fragment of the stem peptide without a bridge (*m/z* = 676) and the same fragment with increasing bridge components (*m/z* = 747, 818, 889), verifying the AAA bridge. A second signature ion is the *m/z* = 431, which represent a stem-fragment with AK plus the entire bridge. In the competent ScrI expression strain (bottom, strain MM481) we see the same stem fragment without bridge (*m/z* = 676), but a shifted series of +30 Da compared to the Δ*scrI* mutant, and it is clear that the + 30 Da shift occurs in position one indicating an Ala→Thr modification. Similarly, the stem-fragment with AK plus the entire bridge is now 30 Da heavier. *m/z* = 555 was determined to be AqKA-[3-NH2-AAA], while *m/z* = 570 was determined to be (AqKA-[3-NH2-TAA]. A: Alanine, q: γ-iso-Glutamine; K: Lysine.

### Supporting tables

**Table S1** Characteristics of *Streptococcus* species tested for sensitivity to ScrM, including interpeptide bridges, cell wall polysaccharides and fratricins and their domains

| Species | Group | Fratricin | Fratricin domains[1] | Interpeptide bridge[2] | Cell wall polysaccharide[3] |
|---|---|---|---|---|---|
| *S. dysgalactiae* | Pyogenic | ScrM | CHAP-SH3b-CCD | $(Ala)_2$-(Ala) | Rha backbone, linked GalNAc |
| *S. pyogenes* | Pyogenic | ScrM | CHAP-SH3b-CCD | $(Ala)_2$-(Ala) | Rha backbone, linked Glc, P, Gly |
| *S. agalactiae* | Pyogenic | Zoocin A | Peptidase_M23 | $(Ala)_2$/Ala-Ser | Rha backbone, linked Rha, Glu, P, Gal, GlcNac |
| *S. pneumoniae* | Mitis | CbpD | CHAP-$(SH3b)_2$-CBD | $(Ala)_2$/Ala-Ser/direct | WTA and LTA |
| *S. mitis* | Mitis | CbpD | CHAP-SH3b-CBD | - | WTA and LTA |
| *S. cristatus* | Mitis | LytF | $(Bsp)_4$-CHAP | - | - |
| *S. mutans* | Mutans | LytF | $(Bsp)_4$-CHAP | Thr-Ala | Rha backbone, linked Glc, P, Gly |
| *S. sobrinus* | Mutans | - | Peptidase-$(Ricin)_2$ | - | - |
| *S. thermophilus* | Salivarius | - | CHAP-CCD | $(Ala)_2$-(Ala) | Rha and Glc backbone, linked Rha, GlcNAc, Gal, Glc |
| *S. vestibularis* | Salivarius | - | CHAP-CCD | - | - |
| *S. suis* | Suis | CrfP | CHAP-SH3b-SH3b | - | - |
| *S. equinus* | Bovis | LytF | $(Bsp)_4$-CHAP | Thr-Ala | - |

[1]CHAP: Cysteine, Histidine-dependent Amidohydrolase/Peptidase; SH3b: Scr Homology 3 domain, bacterial; CCD: conserved C-terminal domain; Bsp: group B streptococcal secreted protein domain. Based on (1).
[2] Based on (2). Ala: Alanine; Ser: Serine; Thr: Threonine
[3] Rha: Rhamnan; Gal: Galactose; Glc: Glucose; P: Phosphate; Gly: Glycerol; Glu: Glucitol; GlcNAc: N-acetylglucosamine; GalNAc: N-acetylgalactosamine; WTA: Wall Teichoic Acid; LTA: Lipoteichoic Acid. Based on (3).

**Table S2** Top 4 predicted reduced muropeptides by PGN_MS2 and MS-DIAL in *S. dysgalactiae* Δ*scrI*.

| Theoretical neutral mass (Da) | Measured neutral mass (Da) | Proposed structure[1] | MS-DIAL score |
|---|---|---|---|
| 569.250 | 569.247 | (GlcNAc)(MurNAc)-A | 1.28 |
| 835.443 | 835.438 | (MurNAc)-AqKA[3-NH2-AA] | 1.14 |
| | | (MurNAc)-AqKAA[3-NH2-A] | 1.06 |
| | | (MurNAc)-AqK[3-NH2-AAA] | 1.06 |
| 906.480 | 906.476 | (MurNAc)-AqKAA[3-NH2-AA] | 1.19 |
| | | (MurNAc)-AqKA[3-NH2-AAA] | 1.07 |
| 967.478 | 967.473 | (GlcNAc)(MurNAc)-AqKAA | 1.23 |
| | | (GlcNAc)(MurNAc)-AqKA[3-NH2-A] | 1.16 |
| | | (GlcNAc)(MurNAc)-AqKA[3-NH2-AA] | 1.04 |
| 1038.523 | 1038.517 | (GlcNAc)(MurNAc)-AqKAA[3-NH2-A] | 1.33 |
| | | (GlcNAc)(MurNAc)-AqKA[3-NH2-AA] | 1.30 |
| | | (GlcNAc)(MurNAc)-AqK[3-NH2-AAA] | 1.15 |
| 1038.523 | 1038.517 | (GlcNAc)(MurNAc)-AqKA[3-NH2-AA] | 1.24 |
| | | (GlcNAc)(MurNAc)-AqKAA[3-NH2-A] | 1.21 |
| | | (GlcNAc)(MurNAc)-AqK[3-NH2-AAA] | 1.11 |
| 1080.533 | 1080.528 | (GlcNAc)(MurNAc-OAc)-AqKA[3-NH2-AA] | 1.19 |
| | | (GlcNAc)(MurNAc-OAc)-AqKAA[3-NH2-A] | 1.17 |
| | | (GlcNAc)(MurNAc-OAc)-AqK[3-NH2-AAA] | 1.06 |
| 1109.560 | 1109.553 | (GlcNAc)(MurNAc)-AqKAA[3-NH2-AA] | 1.27 |
| | | (GlcNAc)(MurNAc)-AqKA[3-NH2-AAA]* | 1.17 |
| | | (GlcNAc)(MurNAc)-AqK[3-NH2-AAAA] | 1.0 |
| 1125.555 | 1125.548 | (GlcNAc)(MurNAc)-AqKAA[3-NH2-SA] | 1.06 |
| | | (GlcNAc)(MurNAc)-AqKAA[3-NH2-AS] | 1.03 |
| | | (GlcNAc)(MurNAc)-AqKAA[3-NH2-TG] | 1.02 |
| | | (GlcNAc)(MurNAc)-AqKAA[3-NH2-GT] | 1.01 |
| 1151.570 | 1151.564 | (GlcNAc)(MurNAc-OAc)-AqKAA[3-NH2-AA] | 1.22 |
| | | (GlcNAc)(MurNAc-OAc)-AqKA[3-NH2-AAA] | 1.13 |
| | | (GlcNAc)(MurNAc-OAc)-AqK[3-NH2-AAAA] | 1.03 |
| 1166.581 | 1166.575 | (GlcNAc)(MurNAc)-AqKAA[3-NH2-AAG] | 1.04 |
| | | (GlcNAc)(MurNAc)-AqKA[3-NH2-AAAG]* | 1.04 |
| | | (GlcNAc)(MurNAc)-AqKAA[3-NH2-AGA] | 1.03 |
| | | (GlcNAc)(MurNAc)-AqKAA[3-NH2-GAA] | 1.02 |
| 1180.597 | 1180.590 | (GlcNAc)(MurNAc)-AqKAA[3-NH2-AAA] | 1.24 |
| | | (GlcNAc)(MurNAc)-AqKA[3-NH2-AAAA] | 1.15 |

[1] GlcNAc: N-acetylglucosamine; MurNAc: N-acetylmuramic acid; OAc: O-acetylation at MurNAc. A: Alanine; q: iso-(γ)-Glutamine; K: Lysine; T: Threonine; G: Glycine. Interpeptide bridges are given in brackets. Asterisk (*) indicate structure confirmed by *de novo* annotation

**Table S3** Top 4 predicted reduced muropeptides by PGN_MS2 and MS-DIAL in *S. dysgalactiae* Δ*scrip*, Ptet-*scrI*.

| Theoretical neutral mass (Da) | Measured neutral mass (Da) | Proposed structure[1] | MS-DIAL score |
|---|---|---|---|
| 697.309 | 697.305 | (GlcNAc)(MurNAc)-Aq | 1.13 |
| 794.409 | 794.405 | (MurNAc)-AqK[3-NH2-TA] | 1.22 |
| | | (MurNAc)-AqK[3-NH2-AT] | 1.13 |
| | | (MurNAc)-AqKA[3-NH2-T] | 1.10 |
| 865.454 | 865.449 | (MurNAc)-AqKAA[3-NH2-T] | 1.16 |
| | | (MurNAc)-AqKA[3-NH2-TA] | 1.11 |
| | | (MurNAc)-AqKA[3-NH2-AT] | 1.09 |
| 906.480 | 906.476 | (MurNAc)-AqKAA[3-NH2-AA] | 1.15 |
| | | (MurNAc)-AqKA[3-NH2-AAA] | 1.04 |
| 926.452 | 926.446 | (GlcNAc)(MurNAc)-AqK[3-NH2-T] | 1.26 |
| 936.491 | 936.486 | (MurNAc)-AqKAA[3-NH2-TA] | 1.18 |
| | | (MurNAc)-AqKAA[3-NH2-AT] | 1.14 |
| | | (MurNAc)-AqKA[3-NH2-ATA] | 1.05 |
| | | (MurNAc)-AqKA[3-NH2-AAT] | 1.00 |
| 967.478 | 967.472 | (GlcNAc)(MurNAc)-AqK[3-NH2-AA] | 1.23 |
| | | (GlcNAc)(MurNAc)-AqKA[3-NH2-A] | 1.15 |
| | | (GlcNAc)(MurNAc)-AqKAA | 1.13 |
| 997.496 | 997.490 | (GlcNAc)(MurNAc)-AqK[3-NH2-TA] | 1.30 |
| | | (GlcNAc)(MurNAc)-AqKA[3-NH2-T] | 1.23 |
| | | (GlcNAc)(MurNAc)-AqK[3-NH2-AT] | 1.23 |
| 1038.523 | 1038.517 | (GlcNAc)(MurNAc)-AqKAA[3-NH2-A] | 1.33 |
| | | (GlcNAc)(MurNAc)-AqKA[3-NH2-AA] | 1.28 |
| | | (GlcNAc)(MurNAc)-AqK[3-NH2-AAA] | 1.13 |
| 1068.533 | 1068.527 | (GlcNAc)(MurNAc)-AqKA[3-NH2-TA] | 1.28 |
| | | (GlcNAc)(MurNAc)-AqKA[3-NH2-AT] | 1.20 |
| | | (GlcNAc)(MurNAc)-AqKAA[3-NH2-T] | 1.20 |
| | | (GlcNAc)(MurNAc)-AqK[3-NH2-ATA] | 1.11 |
| 1080.533 | 1080.530 | (GlcNAc)(MurNAc-OAc)-AqKA[3-NH2-AA] | 1.13 |
| | | (GlcNAc)(MurNAc-OAc)-AqKAA[3-NH2-A] | 1.13 |
| | | (GlcNAc)(MurNAc-OAc)-AqK[3-NH2-AAA] | 1.00 |
| 1095.544 | 1095.538 | (GlcNAc)(MurNAc)-AqKA[3-NH2-GAA] | 1.10 |
| | | (GlcNAc)(MurNAc)-AqKA[3-NH2-AGA] | 1.09 |
| | | (GlcNAc)(MurNAc)-AqKA[3-NH2-AAG] | 1.09 |
| | | (GlcNAc)(MurNAc)-AqKAA[3-NH2-GA] | 1.08 |
| 1109.560 | 1109.554 | (GlcNAc)(MurNAc)-AqKAA[3-NH2-AA] | 1.27 |
| | | (GlcNAc)(MurNAc)-AqKA[3-NH2-AAA] | 1.19 |
| | | (GlcNAc)(MurNAc)-AqK[3-NH2-AAAA] | 1.10 |
| 1110.544 | 1110.538 | (GlcNAc)(MurNAc-OAc)-AqKA[3-NH2-TA] | 1.22 |
| | | (GlcNAc)(MurNAc-OAc)-AqKA[3-NH2-AT] | 1.20 |
| | | (GlcNAc)(MurNAc-OAc)-AqKAA[3-NH2-T] | 1.17 |
| | | (GlcNAc)(MurNAc-OAc)-AqK[3-NH2-ATA] | 1.09 |
| 1139.570 | 1139.565 | (GlcNAc)(MurNAc)-AqKAA[3-NH2-TA] | 1.26 |
| | | (GlcNAc)(MurNAc)-AqKAA[3-NH2-AT] | 1.20 |
| | | (GlcNAc)(MurNAc)-AqKA[3-NH2-ATA] | 1.18 |
| | | (GlcNAc)(MurNAc)-AqKA[3-NH2-TAA]* | 1.18 |
| 1151.570 | 1151.565 | (GlcNAc)(MurNAc-OAc)-AqKAA[3-NH2-AA] | 1.18 |
| | | (GlcNAc)(MurNAc-OAc)-AqKA[3-NH2-AAA] | 1.11 |

| | | | |
|---|---|---|---|
| | | (GlcNAc)(MurNAc-OAc)-AqK[3-NH2-AAAA] | 1.01 |
| 1180.597 | 1180.590 | (GlcNAc)(MurNAc)-AqKAA[3-NH2-AAA] | 1.21 |
| | | (GlcNAc)(MurNAc)-AqKA[3-NH2-AAAA] | 1.11 |
| 1181.581 | 1181.575 | (GlcNAc)(MurNAc-OAc)-AqKAA[3-NH2-TA] | 1.21 |
| | | (GlcNAc)(MurNAc-OAc)-AqKAA[3-NH2-AT] | 1.16 |
| | | (GlcNAc)(MurNAc-OAc)-AqKA[3-NH2-TAA] | 1.10 |
| | | (GlcNAc)(MurNAc-OAc)-AqKA[3-NH2-ATA] | 1.10 |
| 1182.576 | 1182.569 | (GlcNAc)(MurNAc)-AqKA[3-NH2-TAGG] | 1.01 |
| 1196.592 | 1196.585 | (GlcNAc)(MurNAc)-AqKA[3-NH2-TAGA] | 1.06 |
| | | (GlcNAc)(MurNAc)-AqKA[3-NH2-TAAG]* | 1.05 |
| 1210.607 | 1210.601 | (GlcNAc)(MurNAc)-AqKAA[3-NH2-TAA] | 1.12 |
| | | (GlcNAc)(MurNAc)-AqKAA[3-NH2-ATA] | 1.06 |
| | | (GlcNAc)(MurNAc)-AqKA[3-NH2-TAAA] | 1.04 |
| | | (GlcNAc)(MurNAc)-AqKA[3-NH2-ATAA] | 1.04 |
| 1252.618 | 1252.612 | (GlcNAc)(MurNAc-OAc)-AqKAA[3-NH2-TAA] | 1.03 |

[1] GlcNAc: N-acetylglucosamine; MurNAc: N-acetylmuramic acid; OAc: O-acetylation at MurNAc. A: Alanine; q: iso-(γ)-Glutamine; K: Lysine; T: Threonine; G: Glycine. Interpeptide bridges are given in brackets. Asterisk (*) indicate structure confirmed by *de novo* annotation

**Table S4** Top 4 predicted reduced muropeptides by PGN_MS2 and MS-DIAL in *S. dysgalactiae* WT

| Theoretical neutral mass (Da) | Measured neutral mass (Da) | Proposed structure[1] | MS-DIAL score |
|---|---|---|---|
| 611.261 | 611.257 | (GlcNAc)(MurNAc-OAc)-A | 1.10 |
| 835.443 | 835.438 | (MurNAc)-AqKA[3-NH2-AA] | 1.17 |
| | | (MurNAc)-AqKAA[3-NH2-A] | 1.09 |
| | | (MurNAc)-AqK[3-NH2-AAA] | 1.05 |
| 906.480 | 906.475 | (MurNAc)-AqKAA[3-NH2-AA] | 1.20 |
| | | (MurNAc)-AqKA[3-NH2-AAA] | 1.05 |
| 967.486 | 967.480 | (GlcNAc)(MurNAc)-AqKAA | 1.22 |
| | | (GlcNAc)(MurNAc)-AqKA[3-NH2-A] | 1.19 |
| | | (GlcNAc)(MurNAc)-AqK[3-NH2-AA] | 1.11 |
| 997.496 | 997.491 | (GlcNAc)(MurNAc)-AqK[3-NH2-TA] | 1.20 |
| | | (GlcNAc)(MurNAc)-AqK[3-NH2-AT] | 1.19 |
| | | (GlcNAc)(MurNAc)-AqKA[3-NH2-T] | 1.17 |
| 1038.523 | 1038.517 | (GlcNAc)(MurNAc)-AqKAA[3-NH2-A] | 1.34 |
| | | (GlcNAc)(MurNAc)-AqKA[3-NH2-AA] | 1.30 |
| | | (GlcNAc)(MurNAc)-AqK[3-NH2-AAA] | 1.16 |
| 1068.533 | 1068.528 | (GlcNAc)(MurNAc)-AqKA[3-NH2-TA] | 1.27 |
| | | (GlcNAc)(MurNAc)-AqKA[3-NH2-AT] | 1.22 |
| | | (GlcNAc)(MurNAc)-AqKAA[3-NH2-T] | 1.20 |
| | | (GlcNAc)(MurNAc)-AqK[3-NH2-ATA] | 1.12 |
| 1080.533 | 1080.528 | (GlcNAc)(MurNAc-OAc)-AqKA[3-NH2-AA] | 1.26 |
| | | (GlcNAc)(MurNAc-OAc)-AqKAA[3-NH2-A] | 1.23 |
| | | (GlcNAc)(MurNAc-OAc)-AqK[3-NH2-AAA] | 1.13 |
| 1109.560 | 1109.553 | (GlcNAc)(MurNAc)-AqKAA[3-NH2-AA] | 1.28 |
| | | (GlcNAc)(MurNAc)-AqKA[3-NH2-AAA] | 1.21 |
| | | (GlcNAc)(MurNAc)-AqK[3-NH2-AAAA] | 1.12 |
| 1125.555 | 1125.548 | (GlcNAc)(MurNAc)-AqKAA[3-NH2-SA] | 1.08 |
| | | (GlcNAc)(MurNAc)-AqKAA[3-NH2-TG] | 1.05 |
| | | (GlcNAc)(MurNAc)-AqKAA[3-NH2-AS] | 1.05 |
| | | (GlcNAc)(MurNAc)-AqKAA[3-NH2-GT] | 1.04 |
| 1139.570 | 1139.564 | (GlcNAc)(MurNAc)-AqKAA[3-NH2-TA] | 1.22 |
| | | (GlcNAc)(MurNAc)-AqKAA[3-NH2-AT] | 1.18 |
| | | (GlcNAc)(MurNAc)-AqKA[3-NH2-ATA] | 1.14 |
| | | (GlcNAc)(MurNAc)-AqKA[3-NH2-TAA] | 1.14 |
| 1151.570 | 1151.564 | (GlcNAc)(MurNAc-OAc)-AqKAA[3-NH2-AA] | 1.24 |
| | | (GlcNAc)(MurNAc-OAc)-AqKA[3-NH2-AAA] | 1.15 |
| | | (GlcNAc)(MurNAc-OAc)-AqK[3-NH2-AAAA] | 1.04 |
| 1180.597 | 1180.590 | (GlcNAc)(MurNAc)-AqKAA[3-NH2-AAA] | 1.24 |
| | | (GlcNAc)(MurNAc)-AqKA[3-NH2-AAAA] | 1.14 |
| 1181.581 | 1181.574 | (GlcNAc)(MurNAc-OAc)-AqKAA[3-NH2-AT] | 1.09 |
| | | (GlcNAc)(MurNAc-OAc)-AqKAA[3-NH2-TA] | 1.09 |
| | | (GlcNAc)(MurNAc-OAc)-AqKA[3-NH2-AAT] | 1.03 |
| | | (GlcNAc)(MurNAc-OAc)-AqKA[3-NH2-ATA] | 1.02 |

[1] GlcNAc: N-acetylglucosamine; MurNAc: N-acetylmuramic acid; OAc: O-acetylation at MurNAc. A: Alanine; q: iso-(γ)-Glutamine; K: Lysine; T: Threonine; G: Glycine. Interpeptide bridges are given in brackets.

**Table S5** Strains used in this study

| Strain | Genotype and characteristics[1] | Reference |
|---|---|---|
| ***E. coli*** | | |
| DH5α | Cloning host | Lab collection |
| IM08B | Cloning host, DH10B, Δ*dcm*, $P_{help}$-*hsdMS*, $P_{N25}$-*hsdS* | (4) |
| BL21 | Expression host, F-, *ompT*, *hsdS*$_B$ ($r_B^-$ $m_B^-$), *gal*, *dcm* (DE3), pLysS, CmR | Invitrogen |
| MM212 | DH5α, pFD116, spcR | (5) |
| MM291 | DH5α, pFD116-$P_{scrM}$-*luc-gfp*, spcR | This study |
| ATS8 | DH5α, pRSET-*scrM*, ampR | This study |
| ATS10 | BL21, pRSET-*scrM*, ampR | This study |
| ATS36 | DH5α, pRSET-*sf-gfp*-SH3b-CCD | This study |
| MM543 | BL21, pRSET-*sf-gfp*- SH3b-CCD | This study |
| JA9 | DH5α, pRSET-*sf-gfp*-CCD*,* ampR | This study |
| JA10 | BL21, pRSET-*sf-gfp*-CCD*,* ampR | This study |
| JA11 | DH5α, pRSET-*sf-gfp*-CCD$^{,G302A,G303A}$, ampR | This study |
| JA15 | BL21, pRSET-*sf-gfp*-CCD, ampR | This study |
| JA19 | DH5α, pRSET-*scrM*$^{G302A,G303A}$, ampR | This study |
| JA20 | BL21, pRSET-*scrM*$^{G302A,G303A}$, ampR | This study |
| MM479 | IM08B, pFD116-Ptet-*scrI*, spcR | This study |
| ***S. dysgalactiae*** | | |
| Stdys021 | *Streptococcus dysgalactiae* subsp. *dysgalactiae* | (6) |
| MA201 | *Streptococcus dysgalactiae* subsp. *dysgalactiae* | (6) |
| iSDSE-NORM37 | *Streptococcus dysgalactiae* subsp. *equisimilis* | (7) |
| MM420 | Stdys021, pFD116-$P_{scrM}$-*luc-gfp*, spcR | This study |
| MM463 | Stdys021, Δ*scrI::kan*, kanR | This study |
| MM481 | Stdys021, Δ*scrI::kan*, pFD116-$P_{tet}$-*scrI*, kanR, spcR | This study |
| **Other streptococci** | | |
| MK175 | ΔbgaA::P*ssbB-luc-gfp* , Δ*comA::ery* | (8) |
| MM515 | *S. pyogenes* iGAS83 | This study |
| RM22 | *S. agalactiae* NCTC8181 | National Collection of Type Cultures |
| RH425 | *S. pneumoniae* | (9) |
| SPH470 | *S. pneumoniae, ΔcomA, m(sf)GFP-mltG; EryR, SmR* | (10) |
| RM42 | *S. mitis* B6 | Hakenbeck, Brückner, Denapaite and Maurer (11) |
| SA9 | *S. cristatus* NCTC12479 | National Collection of Type Cultures |
| SA58 | *S. mutans* UA159 | American Type Culture Collection |
| SA3 | *S. sobrinus* ATCC27352 | American Type Culture Collection |
| STH2 | *S. thermophilus* BAA-250 | American Type Culture Collection |
| RM38 | *S. vestibularis* NCTC12166 | National Collection of Type Cultures |
| RM36 | *S. suis* 2019-40-11936 | Norwegian Veterinary Institute |
| RM23 | *S. equinus* ATCC9812 | American Type Culture Collection |

[1] ampR = ampicillin resistant, spcR = spectinomycin resistant, kanR = kanamycin resistant

**Table S6** Plasmids used in this study

| Plasmid | Description | Reference |
|---|---|---|
| pFD116 | | (12) |
| pFD116-$P_{scrM}$-*luc-gfp* | | This study |
| pFD116-$P_{tet}$-ScrI | | This study |
| pRSET-A | | Invitrogen |
| pRSET-*scrM* | | This study |
| pRSET-*scrM*$^{G302A,G303A}$ | | This study |
| pRSET-sfGFP-SH3b-CCD | | This study |
| pRSET-sfGFP-CCD | | This study |
| pRSET-sfGFP-CCD$^{G302A,G303A}$ | | This study |

**Table S7** Oligos used in this study

| Oligo name | Sequence (5' – 3') |
|---|---|
| **Primers for construction of pFD116-$P_{scrM}$-*luc-gfp* reporter plasmid** | |
| mm79_PscrM_NheI_F | GTGTCTGCTAGCGGCCACCAGAACCAACAAC |
| mm61_PscrM_over_luc_R | ATGTTTTTGGCGGATCTCATTCATTACCTCCTGTAACTAATTATTC |
| mm55_luc_F | ATGAGATCCGCCAAAAACATAAAG |
| mm57_luc_gfp_SalI_R | TGCTTCGTCGACGAATCTTGCTTGGCAAGGTTC |
| **Primers for amplification and sequencing from pRSET-A** | |
| pRSET(F)[a] | AATACGACTCACTATAGGGAGA |
| pRSET(R)[a] | CTAGTTATTGCTCAGCGGT |
| **Primers for construction of pRSET-*scrM*** | |
| ats1_His_TEV_ScrM_F | TACGCATATGCATCATCATCATCATCATGAGAACCTGTACTTCCAAGGTGAACATACAGGAGTTGTTCATG |
| ats2_ScrM_R | CGTAAAGCTTTTATGGCAATTTAGAGGCTTTATC |
| **Primers for construction of pRSET-*scrM*$^{G302A,G303A}$** | |
| jaa3_scrM_G302_G303_F | GCTGCTGGCGCTACTTCTCTTAACTG |
| jaa4_scrM_G302_G303_R | CAGTTAAGAGAAGTAGCGCCAGCAGCAGCTAGCTCCTCGCTAACTAATAG |
| **Primers for construction of pRSET-*sf-gfp-scrM*$^{\Delta CHAP}$** | |
| ats3_His_sfgfp_F | TACGCATATGCATCATCATCATCATCATAAACATCTTACCGGTTCTAAAG |
| ats4_sfGFP_R | TGCGGCCGCTCCACTAGTTTTG |
| ats5_SH3b_over_sfGFP_F | CAAAACTAGTGGAGCGGCCGCACCTAGTGGTGAAGCAAGTAAG |
| ats2_scrM_R | CGTAAAGCTTTTATGGCAATTTAGAGGCTTTATC |
| **Primers for construction of pRSET-*sf-gfp-scrM*$^{\Delta CHAP\Delta SH3B}$ and pRSET-*sf-gfp-scrM*$^{\Delta CHAP\Delta SH3B,G302A,G303A}$** | |
| jaa1_cons_domain_F | AAAAAGACGGGGCAAAAAACAC |
| jaa2_sfgfp_over_cons_domain_R | GTGTTTTTTGCCCCGTCTTTTTACTGCTTCTTTAGAAACCTGTTG |
| **Primers for construction of Δ*scrI::kan*** | |
| mm107_up_scrI_F | GAGGTGATTAATCATAGCCTTC |
| mm109_up_scrI_over_kan_R | CACATTATCCATTAAAAATCAAACTTGCCTCTCTCATCTTCTAAATG |
| mm110_down_scrI_over_kan_F | GTCCAAAAGCATAAGGAAAGAAAAGAGCTCGCCAAAGTATG |
| mm111_down_scrI_R | GGCCTTCCTGAAGACATGC |
| **Primers for amplification of kanamycin resistance cassette** | |
| kan484F[b] | GTTTGATTTTTAATGGATAATGTG |
| rpsL41R[b] | CTTTCCTTATGCTTTTGGAC |
| **Primers for construction of pFD116-Ptet-*scrI*** | |
| mm120_pFD116_R | TTTTGCCTCCTATATGCCTC |
| mm121_pFD116_F | AGATCTGTCCATACCCATGG |
| mm122_scrI_over_pFD116_F | GAGGCATATAGGAGGCAAAAATGACAAAGGTAACATTTTATTC |
| mm123_scrI_over_pFD116_R | CCATGGGTATGGACAGATCTTTAATGCTTATGTTTTAACCG |

[a] Invitrogen

[b] Johnsborg, Eldholm, Bjørnstad and Håvarstein (13)

### Supporting text

**Text S1.** DNA sequences encoding *scrM* and *scrI*.

**>ScrM_encoding_MA021**
ATGAAAAAAATTCATCAATTGTTAGTGTCAGGAGCAATCCTTTTGAGTGTTAATGGTGCTGTATCTTC
AGTTGCATCCACTTTGAATGCAGAGCATACAGGAGTTGTTCATGCAGCTGTCCTTGGGGATAATTATC
CTAGCAAATGGAAAAAGGGATCTGGAATTGATTCTTGGAATATGTATGTTCGTCAGTGCACATCATTT
GTCGCTTTCCGTCTGAGTTCAGCAAATGGTTTTCAGTTGCCCAAAGGCTATGGGAATGCCTGCACTT
GGGGGCATATTGCAAAAAAACAAGGCTATACTGTCAATAAGACCCCTAAAGTCGGGGCAGTGGCGT
GGTTTGATACTAACGCTTTCCAATCTCATGCAACGTATGGTCATGTGGCTTGGGTAGCCGAAGTACGT
GGAGATTCTGTTGTGATTGAGGAATATAATTACAATGCTGGTCAAGGACCTGAGAAATACCATAAGC
GTCAAATCCCCAAAAACCATGTGAGCGGTTATATTCATTTTAAAGATTTGCCTAGTGGTGAAGCAAG
TAAGTCTCAAACAAAAGAACAACAGGTTTCTAAAGAAGCAGTAAAACAAGGAGGAACTTACCATT
TTACTGAGCGTACTCCTGTTAAAGCACAGGCTCAACTCACTAGTCCTGACTTAGCTTATTATAATCCT
GGACAATCTGTTCATTACGATCAAGCTATGACCGTTGACGGTCATGAATGGATTAGCTATCTCAGTTT
TTCAGGAAGTCGACGTTATATCCCAATTAAAAAGACGGAGCAAAAAACACAACAAGTCTCTGAGAC
AACATCGCCTATCAATATTGGAGATAGAGTGACTTTCCCTGGCGTTTTCCGTGTGGATCGTATTGTAA
ACAATCTATTAGTTAGCGAGGAGCTAGCTGGTGGGGGCGCTACTTCTCTTAACTGGATTGATCCCTCA
CCCTTGGATGAAACAGATCGTAAAGGAGTAAAAGCAGGAAATCAAATTTTACAGGCCGGTGAGTTT
TTTGTTATCCCAGGTAACTATAGGGTACTGAAAGTCGACCGACCAAGCAATGGGATTTATGTCAAGA
TTGGATCACGTGGAACATGGTTAACTGCTGATAAAGCCTCTAAATTGCCATAA

**>ScrI_encoding_Stdys021**
ATGACAAAGGTAACATTTTATTCAAAAATAGGAATTTCGGCAGAAGAACACGATGCTTTTGTAACGC
AACATGAGCAAGTCAATTTGTTACAGAGTAGTAATTGGGCTAAGGTTAAAGACCAATGGGAAAATG
AACGGATTGGTATCTACAAAGGAAATCAGCAGGTTGCTTCTTTATCTCTGTTAATTAAACCATTACCT
CTCGGGATGACTATTATCTATATCCCTAGAGGACCAGTCATGGATTATGGGGATTATGATTTGGTAACT
TTTACGATGAACACACTCAAAGATTACGGTAAATTGAAAAAAGCTTTATTTATTAAATGCGATCCTGC
GATACTTTTAAAACAATATTCGCTAGGTCAAGAAGGAGAGAAAAAAACAACTGCTTTAACAGCTATT
GAGAATCTGAAGAAGGCAGGTGCTCATTGGACAGGTCTGACGACAGCTATTGCAGATAGCATCCAA
CCCCGGTTTCAAGCTAATGTTTATCCTGAGAAGGAGCATCACCTCACCTTTCCAAAACACACTAGGC
GTCTGATGAAAGATGCTATGCAGCGTGGGGTAACAACTTATCGTGCAACGCCATCAGAGATTGAACA
GTTTTCAGCAGTTGTTTCATTAACAGAAAAACGAAAAAATATTTCCCTACGTAATAAAGCTTATTTTA
AAAAGTTAATGGCAATCTATGGTGATAGAGCTTACTTACATTTGGCCAAAGTCAACATTTCACAGCA
ATTGACACACTATCAGCAACAATTAGCAGTTGTTAATGAGGAGATTGAACTTACTCAACCACATCAA
AAAAAACGACTCAAGAAATTAGAAGAGCAGAAAAGTTCTTTGGAACGTTACATTGTTGACTTTGAA
ACTTTTGGGACAAGCTCACCTGAAGATGTTGTTATAGCAGGGATTCTTTCCATTTCTCATGGAAATGT
CATGGAGATGCTTTATGCTGGAATGAATGAGGCGTTTAAAAAATGCTATCCGCAATACCTTCTCTATC
CTAAAGTCTTTCAAGATGCTTATCAAGATGGGATTATTTGGGCGAATATGGGAGGTGTAGAAGGAAC
GCTTGACGACGGATTAACAAGGTTCAAGTCTCATTTCTCTCCTGTTATCGAAGAATTTATAGGAGAGT
TTACACTTCCTGTGAGTCCACTGTATGTCCTTGCTAATATGCTTTACACCTTCCGGAAACGGTTAAAA
CATAAGCATTAA

#### References


1. K. H. Berg, T. J. Biørnstad, O. Johnsborg, L. S. Håvarstein, Properties and biological role of streptococcal fratricins. *Appl Environ Microbiol* **78**, 3515-3522 (2012).
2. K. H. Schleifer, O. Kandler, Peptidoglycan types of bacterial cell walls and their taxonomic implications. *Bacteriol Rev* **36**, 407-477 (1972).
3. H. Guérin, S. Kulakauskas, M. P. Chapot-Chartier, Structural variations and roles of rhamnose-rich cell wall polysaccharides in Gram-positive bacteria. *J Biol Chem* **298**, 102488 (2022).
4. I. R. Monk, J. J. Tree, B. P. Howden, T. P. Stinear, T. J. Foster, Complete Bypass of Restriction Systems for Major *Staphylococcus aureus* Lineages. *mBio* **6**, e00308-00315 (2015).
5. M. T. Mårli, O. Oppegaard, D. Porcellato, D. Straume, M. Kjos, Genetic modification of *Streptococcus dysgalactiae* by natural transformation. *mSphere* **0**, e00214-00224 (2024).
6. D. Porcellato *et al.*, Whole genome sequencing reveals possible host species adaptation of *Streptococcus dysgalactiae*. *Sci Rep* **11**, 17350 (2021).
7. A. Kaci *et al.*, Genomic epidemiology of *Streptococcus dysgalactiae* subsp. *equisimilis* strains causing invasive disease in Norway during 2018. *Front Microbiol* **14**, 1171913 (2023).
8. S. Moreno-Gámez *et al.*, Quorum sensing integrates environmental cues, cell density and cell history to control bacterial competence. *Nat Commun* **8**, 854 (2017).
9. O. Johnsborg, L. S. Håvarstein, Pneumococcal LytR, a protein from the LytR-CpsA-Psr family, is essential for normal septum formation in *Streptococcus pneumoniae*. *J Bacteriol* **191**, 5859-5864 (2009).
10. G. A. Stamsås *et al.*, Identification of EloR (Spr1851) as a regulator of cell elongation in *Streptococcus pneumoniae*. *Mol Microbiol* **105**, 954-967 (2017).
11. R. Hakenbeck, R. Brückner, D. Denapaite, P. Maurer, Molecular mechanisms of β-lactam resistance in *Streptococcus pneumoniae*. *Future Microbiol* **7**, 395-410 (2012).
12. F. Depardieu, D. Bikard, Gene silencing with CRISPRi in bacteria and optimization of dCas9 expression levels. *Methods* **172**, 61-75 (2020).
13. O. Johnsborg, V. Eldholm, M. L. Bjørnstad, L. S. Håvarstein, A predatory mechanism dramatically increases the efficiency of lateral gene transfer in *Streptococcus pneumoniae* and related commensal species. *Molecular Microbiology* **69**, 245-253 (2008).
